# Focused ultrasound ablation of melanoma with boiling histotripsy yields abscopal tumor control and antigen-dependent dendritic cell activation

**DOI:** 10.1101/2023.09.02.552844

**Authors:** Eric A. Thim, Lydia E. Kitelinger, Fátima Rivera-Escalera, Alexander S. Mathew, Michael R. Elliott, Timothy N. J. Bullock, Richard J. Price

**Author notes:** These authors contributed equally: E. Andrew Thim and Lydia E. Kitelinger. Corresponding Authors: Richard J. Price, Ph.D. Department of Biomedical Engineering Box 800759, Health System University of Virginia Charlottesville, VA 22908, USA; Timothy N. J. Bullock, Ph.D. Department of Pathology Box 801386, Health System University of Virginia Charlottesville, VA 22908, USA.

## Abstract

**Background:** Boiling histotripsy (BH), a mechanical focused ultrasound ablation strategy, can elicit intriguing signatures of anti-tumor immunity. However, the influence of BH on dendritic cell function is unknown, compromising our ability to optimally combine BH with immunotherapies to control metastatic disease.

**Methods:** BH was applied using a sparse scan (1 mm spacing between sonications) protocol to B16F10-ZsGreen melanoma in bilateral and unilateral settings. Ipsilateral and contralateral tumor growth was measured. Flow cytometry was used to track ZsGreen antigen and assess how BH drives dendritic cell behavior.

**Results:** BH monotherapy elicited ipsilateral and abscopal tumor control in this highly aggressive model. Tumor antigen presence in immune cells in the tumor-draining lymph nodes (TDLNs) was ∼3-fold greater at 24h after BH, but this abated by 96h. B cells, macrophages, monocytes, granulocytes, and both conventional dendritic cell subsets (i.e. cDC1s and cDC2s) acquired markedly more antigen with BH. BH drove activation of both cDC subsets, with activation being dependent upon tumor antigen acquisition. Our data also suggest that BH-liberated tumor antigen is complexed with damage-associated molecular patterns (DAMPs) and that cDCs do not traffic to the TDLN with antigen. Rather, they acquire antigen as it flows through afferent lymph vessels into the TDLN.

**Conclusion:** When applied with a sparse scan protocol, BH monotherapy elicits abscopal melanoma control and shapes dendritic cell function through several previously unappreciated mechanisms. These results offer new insight into how to best combine BH with immunotherapies for the treatment of metastatic melanoma.

**Graphical Abstract:** 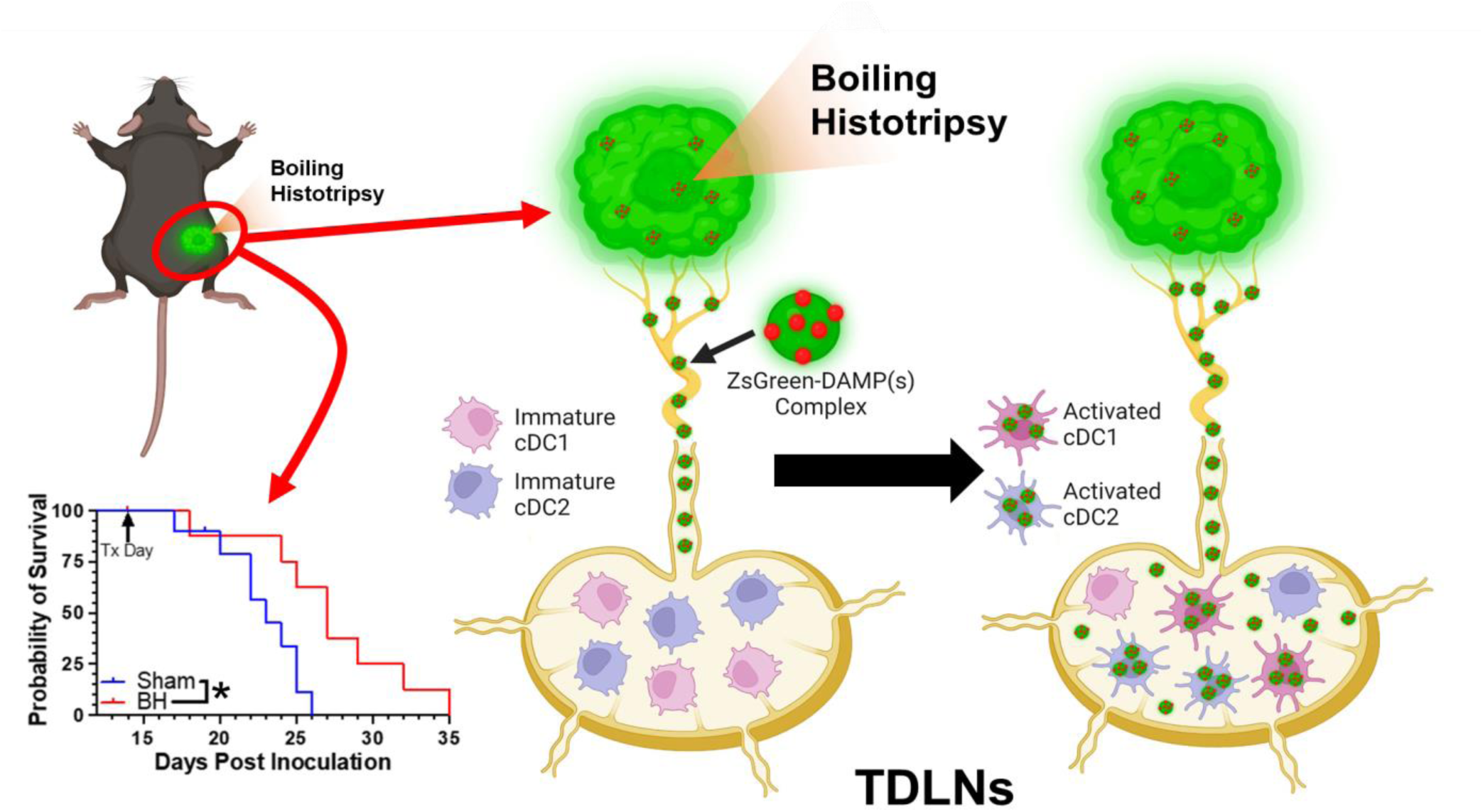

## Introduction

Melanoma diagnoses continue to rise, with ∼105,000 new cases predicted for 2023. Despite significant recent advances in treatment with targeted therapies and immunotherapies, melanoma patients who experience distant metastatic spread still have only a 32% 5-year survival rate [1]. Immunotherapies aimed at increasing the endogenous immune response against melanoma are now standard in the clinical armamentarium. Such immunotherapies have a variety of targets, including programmed death receptor/ligand-1 (PD-1/PD-L1), cytotoxic T-lymphocyte associated protein 4 (CTLA-4), and interleukin 2 (IL-2) [2,3]. However, many patients still do not experience the survival benefits these therapies can offer. Their tumors, which are collectively termed immunologically “cold” [2,4], typically have a paucity of T lymphocyte (T cell) infiltration. Limited T cell presence within tumors can be the result of inadequate tumor antigen acquisition and presentation by dendritic cells (DC), which serve as obligate activators of tumor-specific T cells; a failure of DC to traffic to lymph nodes to interact with the T cell repertoire; a surplus of immunosuppressive cells (e.g. regulator T cells [Tregs] and myeloid derived suppressor cells [MDSCs]) that suppress T and DC activity; and/or an inability of activated T cells to traffic to and persist in tumors [2,4]. There is a clear need for a treatment modality that can transform a “cold tumor” into a “hot tumor” for increased responses to immunotherapies.

Focused ultrasound (FUS), a term referring to the concentration of acoustic energy into a small focus to create bioeffects in tissue, holds considerable promise as a minimally-invasive means for transforming “cold” tumors into “hot” tumors, while limiting off-target and side effects. FUS is a versatile treatment modality that is not limited by dose and may be repeated often due to its non-ionizing nature [5–7]. Tumor tissue fragmentation may be achieved through a specific form of FUS known as histotripsy, wherein short-duration pulses at high intensity elicit mechanical disintegration through generation and subsequent manipulation of vapor bubble activity [8]. While multiple forms of histotripsy exist [8], the current study is centered on so-called “boiling histotripsy” (BH), wherein high pressure, millisecond long, FUS pulses are deployed. BH beneficially modulates immune landscape [9–16] and cooperates with immunological checkpoint inhibitors to control tumor growth [13–16]. Of particular note, BH has been reported to modulate dendritic cell activation and migration [11,12,16], repolarize tumor-associate macrophages [13,15], and enhance T cell representation in tumors [11,13–16]. Pre-clinically, BH has also been combined with αCTLA4 and αPD1 to treat neuroblastoma[14], αPD-L1 to treat triple negative and HER2 breast tumors[13], αCD40 agonist to treat melanoma[15], and αPD-1 to treat 4T1 breast tumors[16].

However, despite the clear potential for BH to stimulate adaptive immune responses against solid tumors, there are still important gaps in our understanding of how BH affects key elements of the cancer-immunity cycle. Until these knowledge gaps are filled, our ability to optimally combine immunotherapies with BH to drive systemic anti-tumor immune responses will be compromised. In particular, many of these gaps center on how and where BH affects tumor antigen trafficking and acquisition. For example, BH-driven tumor antigen trafficking to TDLNs has not been directly measured and we don’t know which antigen presenting cell (APC) type actually acquire antigen in TDLNs. Such knowledge will be invaluable for better defining the time course of administration of immunotherapies intended to synergize with BH via augmented tumor antigen acquisition by APCs, as well as for designing studies aimed at defining how BH modulates phagocytic activity of APCs. Furthermore, it is unknown as to whether/how BH-liberated tumor antigen is partitioned amongst DC subsets (i.e., cDC1 vs. cDC2). Because CD8^+^ T cells are generally thought to be primarily activated by cDC1s [17–19], while CD4^+^ T cells require cDC2s for initial priming [17], defining both the activation and relative acquisition of tumor antigen by each DC subset may help identify opportunities for therapeutically tuning the relative contributions of effector and helper T cells to the BH-induced anti-tumor immunity. Moreover, while DC maturation has been reported in response to BH, it unknown whether activation depends upon acquisition of BH-liberated tumor antigen in vivo and whether tumor antigen is preferentially acquired by DCs in the tumor microenvironment or in the TDLN. Such knowledge will inform proper tuning of the intensity and volumetric fraction of BH to optimally elicit anti-tumor immunity. For example, if DC activation depends on tumor antigen acquisition, a more aggressive liberation of antigen by BH would be warranted. On the other hand, evidence for DC acquisition of antigen in the tumor microenvironment could suggest that reducing the volumetric fraction of BH treatment could improve anti-tumor immunity by sparing intratumoral DCs from ablation.

Here, we directly address these key gaps in our understanding of how BH drives anti-tumor immunity in a mouse model of melanoma. By employing a B16F10 cell line that stably expresses ZsGreen (ZsG) (i.e., B16F10-ZsG) as a model tumor antigen in combination with a BH treatment scheme that elicits abscopal tumor control, we specifically investigated (i) tumor BH-induced antigen drainage to lymph nodes, (ii) tumor antigen acquisition and partitioning by phagocytic immune cells and DC subsets in the TDLN, (iii) DC activation as a function of tumor antigen acquisition, and (iv) the trafficking potential of antigen positive DCs through CD8α^+^ (tissue resident) and CD103^+^ (migratory) cDC1 subpopulations. Our findings yield new insights into DC behavior in the setting of abscopal tumor control, while also providing guidance for how immunotherapeutic manipulations may be rationally combined with BH to further control of systemic disease.

## Results

### Boiling Histotripsy Elicits Primary and Abscopal Tumor Control

We first developed a BH treatment protocol that yields abscopal control of distal disease for a melanoma model (B16F10-ZsG) that is stably-transfected to express a fluorescent protein (ZsG) in cytoplasm[20]. It has been shown that ZsG persists in intracellular compartments, allowing for tracking of this fluorescent protein in APCs by flow cytometry[21]. Because the B16F10-ZsG melanoma model was deployed for all experiments in this study, it is henceforth often referred to as “tumor” or “melanoma.” “Primary” refers to ipsilateral (treated) while “secondary” refers to contralateral (untreated). The primary tumors were chosen for treatment as the larger of the two tumors. When applied to the ipsilateral tumor in a bilateral setting at 14 days post-inoculation, our BH regimen (Figure 1A; FUS parameters provided in Figure S1) significantly improved survival (Figure 1B) and controlled ipsilateral tumors (Figure 1C-E), with the “area under the curve” (AUC) metric showing a highly significant ∼40% reduction in integrated tumor burden (Figure 1E). For contralateral tumors not directly exposed to BH, multi-variate statistical analysis of the modeled growth curves showed a strong trend toward growth control (Figure 1F-G), with the AUC metric (Figure 1H) yielding a significant reduction in integrated contralateral tumor burden. In all, this indicates that our BH monotherapy protocol elicits abscopal tumor control in this model.

**Figure 1.**
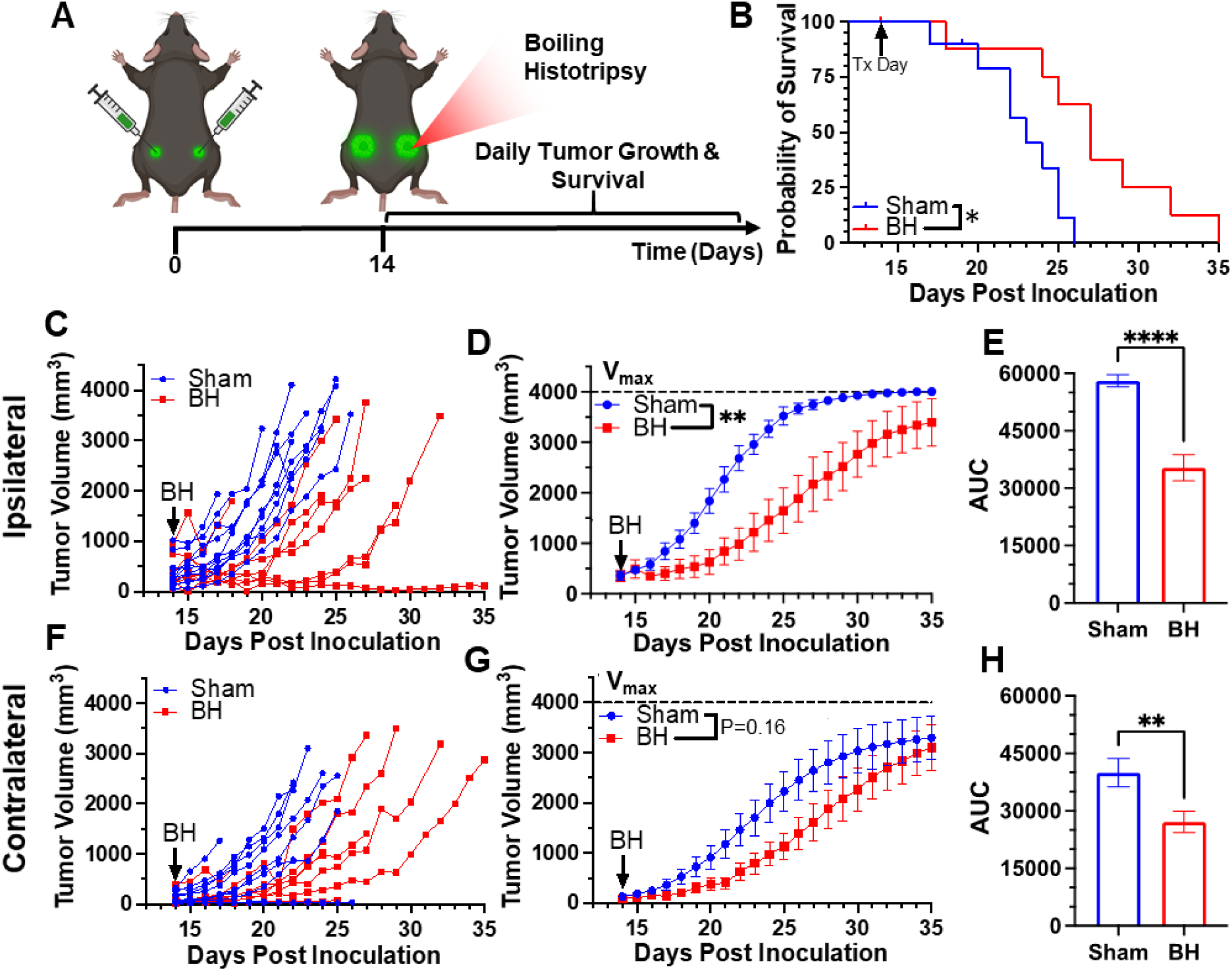
BH yields primary and abscopal control of B16F10-ZsG melanoma. 4×10^5^ B16F10-ZsG cells were inoculated in the left and right flanks of C57/Bl6 mice and tumors were exposed to BH or sham treatment 14 days post inoculation. **A.** Timeline for inoculation and treatment. **B.** Kaplan-Meier curve depicting overall survival (significance assessed by log-rank (Mantel-Cox) test: ∗ P<0.05). **C-H.** Ipsilateral and contralateral tumor growth. **C & F.** Individual tumor growth curves. **D & G.** Average logistic modeled tumor growth. n=8-11 per group. Full model, two-way repeated measures ANOVA from day 14 to 35, fixed effects: ∗∗ P<0.01. **E & H.** Area under the logistic average curve (AUC). Unpaired Welch’s t-test: ∗∗ p < 0.01, ∗∗∗∗ p < 0.0001. Means ± SEM.

### Boiling Histotripsy Transiently Increases Tumor Antigen Acquisition by Immune Cells

After establishing that this BH treatment regimen controls distal tumor growth (Figure 1), we examined the time course of antigen acquisition by all immune cells, as identified by CD45^+^ staining, in the TDLNs in a unilateral B16F10-ZsG model in response to BH (Figure 2A). The ZsG fluorescent antigen allowed for the tracking of antigen in TDLN cells (Figure 2B; gating strategy provided in Figure S2). We found that BH elicited a nearly three-fold increase in the proportion of ZsG^+^ CD45^+^ immune cells 24 h post-treatment (Figure 2C). However, by 96 h, ZsG antigen presence in CD45^+^ cells returned to near baseline levels. Interestingly, the contralateral non-TDLN (CLN) exhibited low-levels of baseline ZsG antigen (Figure S3), with the proportion of CD45^+^ cells that are ZsG^+^ being ∼15-fold lower than those in the baseline ipsilateral TDLN. Nonetheless, BH did not alter contralateral tumor antigen presence, indicating that BH does not increase circulating tumor antigen. Further, BH did not increase the number or proportion of CD45^-^ZsG^+^ cells either in the CLN or TDLN (Figure S4), suggesting that BH did not promote dissemination of tumors cells to lymph nodes, mitigating the concern that mechanically destroying tumors could increase the release of tumor cells to distant sites (e.g., TDLNs) [22–24].

**Figure 2.**
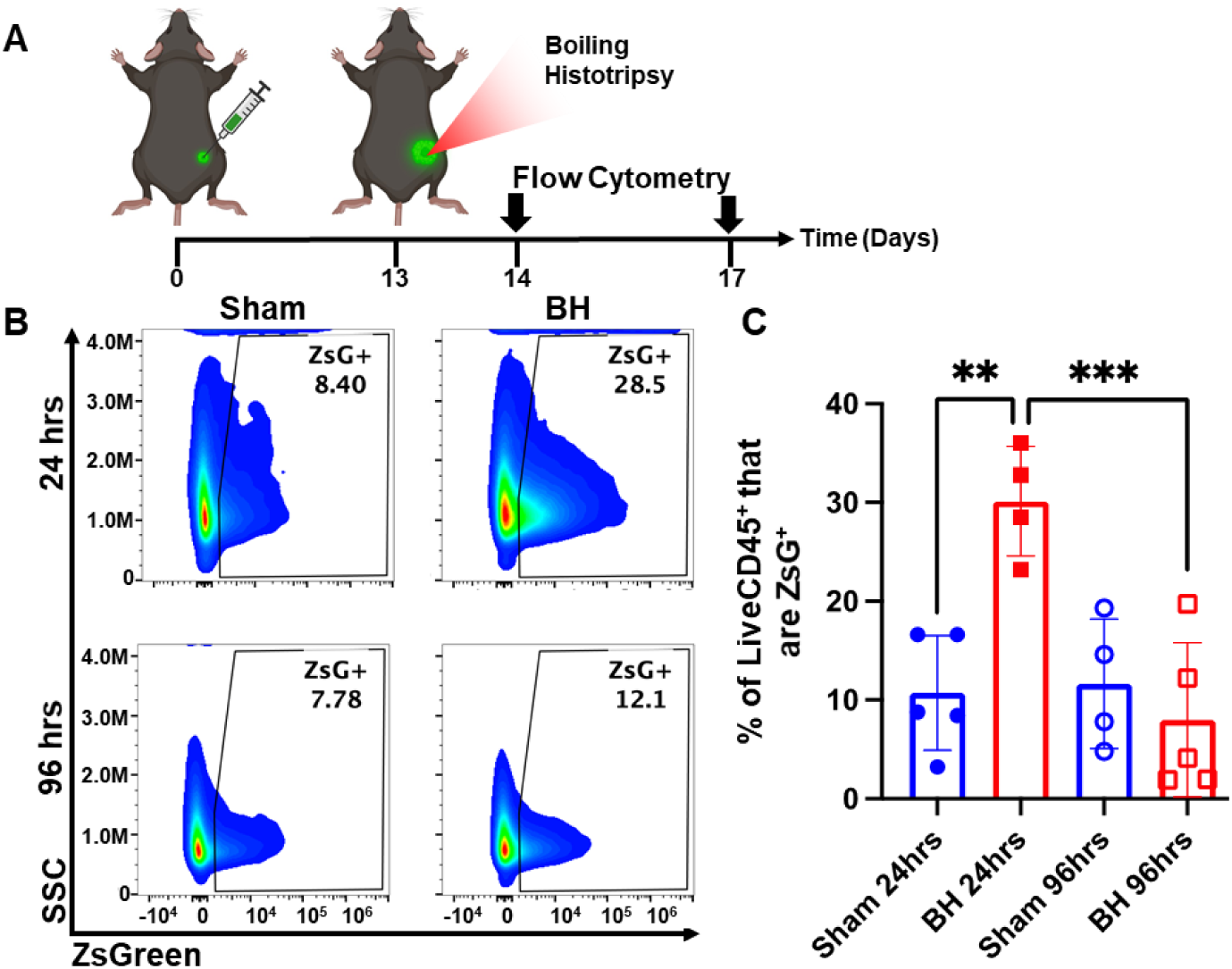
BH transiently increases tumor antigen acquisition by immune cells in the tumor draining lymph nodes. 4×10^5^ B16F10-ZsG cells were inoculated in the right flanks of C57/Bl6 mice and tumors were exposed to BH or sham treatment 13 days post inoculation. **A.** Timeline for inoculation, treatment, and harvest for flow cytometry. Inguinal, axial, and brachial lymph nodes were harvested and pooled 24 h and 96 h post treatment. **B.** Scatter density plots indicating percent of LiveCD45^+^ cells that are ZsG^+^. **C.** Bar graph of flow cytometry analysis data. n=4-5 per group. Full model, two-way ANOVA followed by Tukey multiple comparison test: ∗∗ p < 0.01, ∗∗∗ p < 0.001. Means ± SEM.

### Boiling Histotripsy Induces Antigen Acquisition by Multiple Phagocytic Cell Types

We next asked which immune cell types in the TDLN acquired tumor antigen at 24 h after BH, as antigen partitioning after BH is currently unknown and could significantly impact anti-tumor immunity (Figure 3A). We specifically examined ZsG acquisition by antigen presenting and phagocytic cells such as DCs, B cells, macrophages, monocytes, and granulocytes (gating strategy provided in Figure S5.) For all APC and phagocytic cell types examined, we observed an increase in both the number (Figure 3B-F) and proportion (Figure 3G-K) of ZsG^+^ cells 24 h post BH treatment. Further, the amount of ZsG antigen each cell type acquired, as quantified by geometric mean fluorescent (GMF) intensity (Figure 3L-P), increased in DCs (2-fold), macrophages (3.5-fold) and granulocytes (2.5-fold) as a result of BH treatment. Interestingly, despite B cells exhibiting the greatest increase in both the number and proportion of cells acquiring ZsG, no difference in the GMF of ZsG was observed. Because all cell types acquired ZsG in Figure 3, it is important to highlight the negative control experiment for ZsG acquisition. To this end, we examined ZsG positivity of a non-APC and non-phagocytic cell type (i.e., CD8^+^ T cells; Figure S6A). As expected, we found an extremely low proportion of CD8^+^ T cells acquired ZsG. Further, BH did not change this proportion (Figure S6B). Altogether, this analysis shows that BH enhanced ZsG tumor antigen presence in all examined phagocytic and antigen presenting cell types in TDLNs 24 h post treatment.

**Figure 3.**
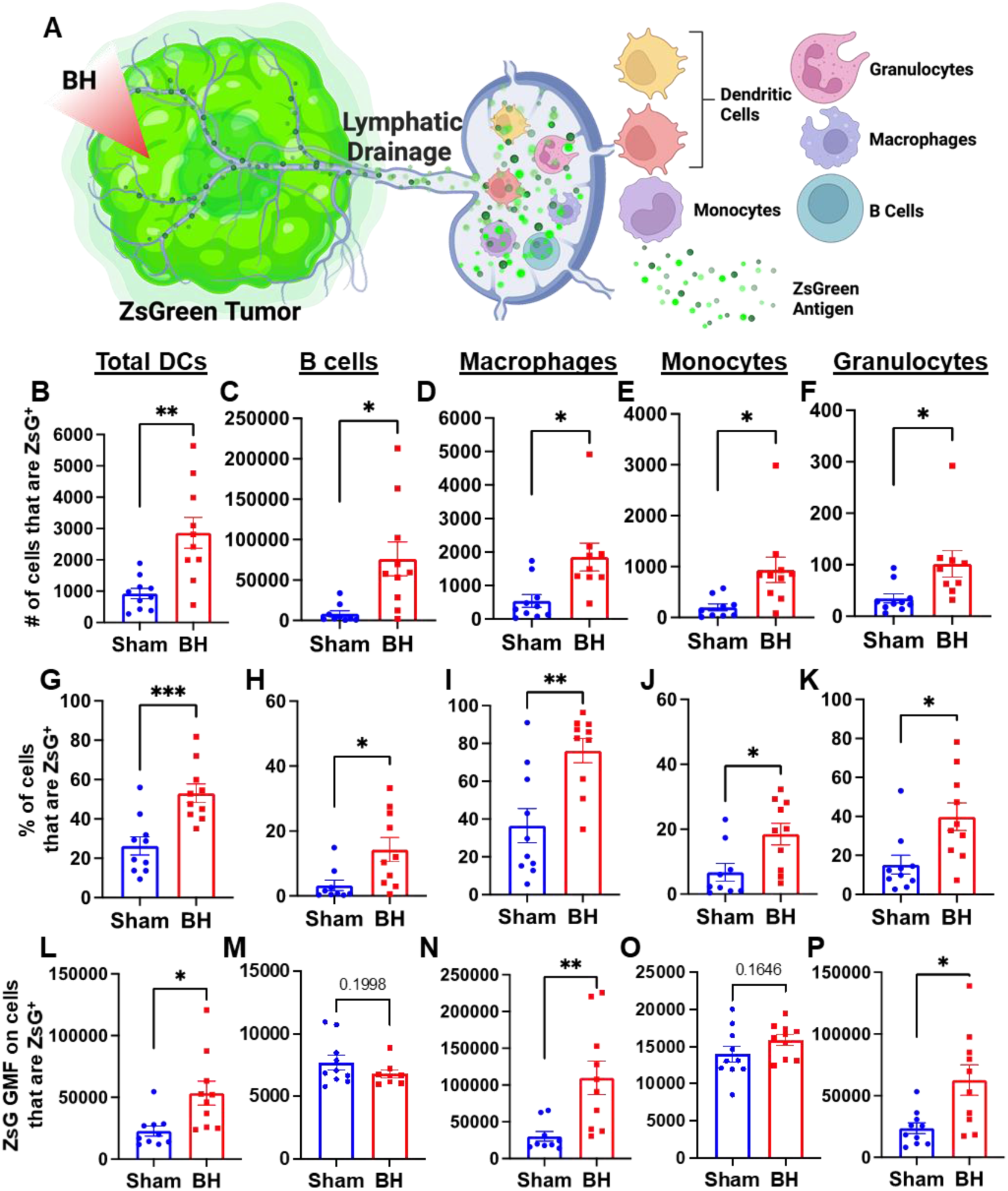
BH enhances tumor antigen acquisition by antigen presenting and phagocytic cells. **A.** Diagram outlining immune cells interrogated for ZsG positivity. **B-F.** Bar graphs of ZsG^+^ cell counts. **G-K.** Bar graphs of proportions of ZsG^+^ cells. **L-P.** Bar graphs of geometric mean fluorescent intensity (GMF) of ZsG on ZsG^+^ cells. **B, G, L.** Total DCs (CD11c^+^MHCII^+^). **C, H, M.** B cells (CD19^+^CD3^-^). **D, I, N.** Macrophages (CD11b^+^F4/80^+^). **E, J, O.** Monocytes (CD11b^+^F4/80^-^Ly6C^+^Ly6G^-^). **F, K, P.** Granulocytes (CD11b^+^F4/80^-^Ly6C^mid^Ly6G^+^). n = 10 per group. Unpaired Welch’s t-test: ∗ p < 0.05, ∗∗ p < 0.01, ∗∗∗ p < 0.001. Means ± SEM.

### Conventional DCs Acquire Antigen in Response to Boiling Histotripsy

We observed an almost 3-fold increase in the number and proportion of DCs that acquired ZsG after BH (Figure 3B and G). DC subsets (i.e., cDC1 and cDC2) have distinct phenotypes and functions that can differentially affect anti-tumor immune responses (Figure 4A) (cDC1: CD8^+^ T cell activation and CD4^+^ T cell licensing; cDC2: CD4^+^ T cell priming). Thus, to understand whether DCs differentially acquire BH-liberated tumor antigen, we separated cDC1s (XCR1^+^) and cDC2s (XCR1^-^CD11b^+^SIRPα^+^) from the total DC population (gating strategy provided in Figure S7) and measured changes in ZsG expression (Figure 4B) for cDC1s (Figure 4B; Top) and cDC2s (Figure 4B; Bottom) 24 h post-treatment. When quantified, the number and proportion of both cDC1s (Figure 4C and D) and cDC2s (Figure 4F and G) that are ZsG^+^ increased significantly with BH. When examining GMF, the amount of ZsG per cell on cDC2s significantly increased with BH (Figure 4H), while the amount of ZsG per cell on cDC1s trended toward an increase (Figure 4E). This shows that within the DC compartment, BH enhanced tumor antigen expression by both cDC1s and cDC2s.

**Figure 4.**
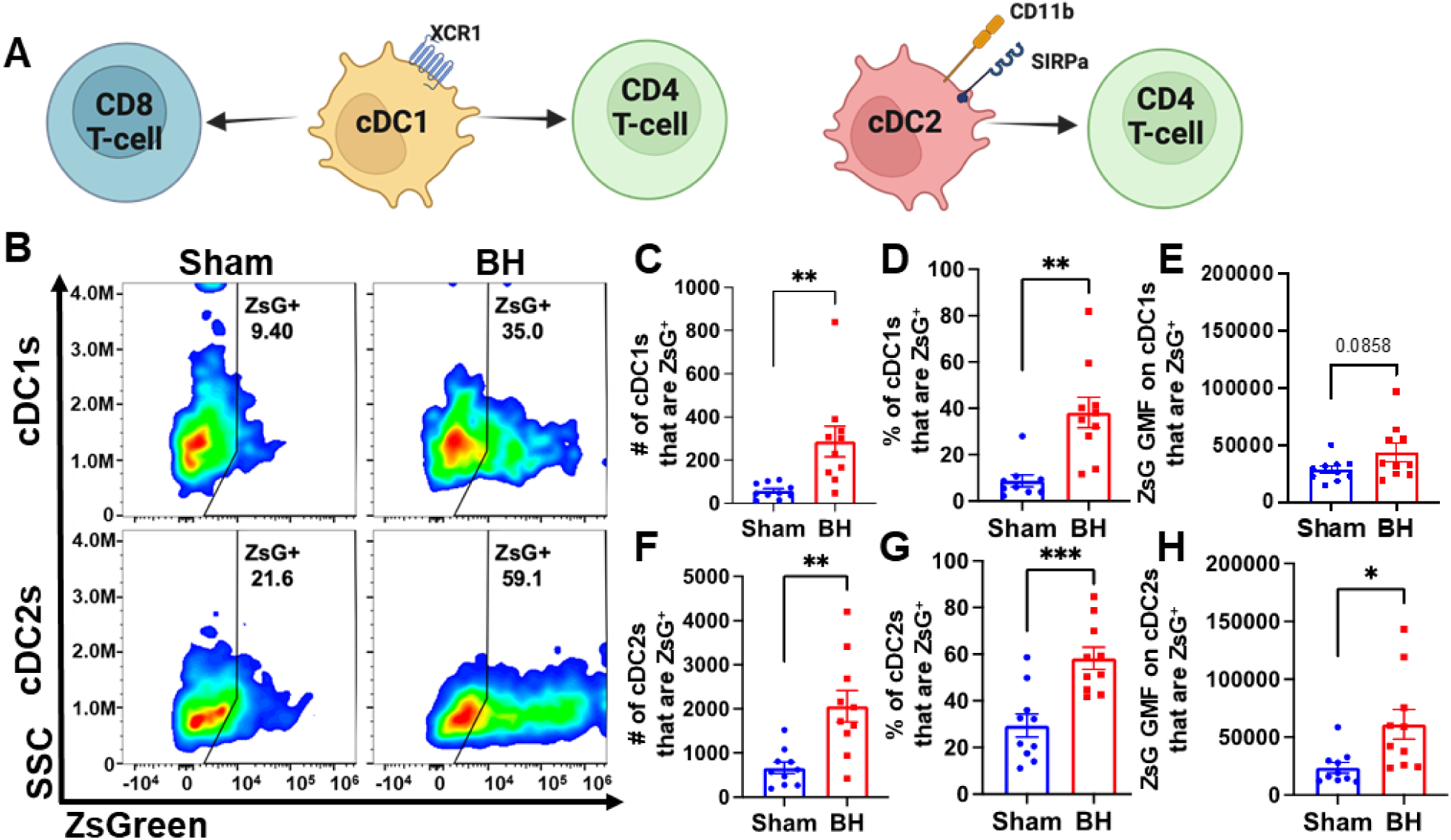
BH increases ZsG tumor antigen presence in cDCs in the tumor draining lymph nodes. **A.** Overview of CD8^+^ and CD4^+^ T cell activation by cDC1s and cDC2s. **B.** Density scatter plots of side scatter vs ZsG for cDC1s (top) and cDC2s (bottom). **C & F.** Bar graphs of numbers of cDCs that are ZsG^+^. **D & G.** Bar graphs of percentages of cDCs that are ZsG^+^. **E & H.** Bar graphs of ZsG GMF on cDCs that are ZsG^+^. n=10 per group. Unpaired Welch’s t-test: ∗ p < 0.05, ∗∗ p < 0.01, ∗∗∗ p < 0.001. Means ± SEM.

### Boiling Histotripsy Activates cDCs

Knowing that both cDC1s and cDC2s exhibit enhanced tumor antigen acquisition after BH, we next asked whether BH elicited changes in the activation status of cDCs present within the TDLNs 24 h post treatment. cDC1 and/or cDC2 activation, which has not been previously reported or characterized in response to BH, is an essential step in the cancer immunity cycle as it is required for effective T cell priming and activation to elicit anti-tumor immunity. As CD86 is a co-stimulatory molecule upregulated on the surface of DCs as they undergo activation and maturation (Figure 5A), we first examined CD86 presence on the surface of cDC1s and cDC2s (Figure 5B). We found that BH stimulates greater overall CD86 expression per cell in both cDC subsets (Figure 5C and F). Using the flow cytometry gating strategy described in Figure S7, we identified a secondary CD86^hi^MHCII^hi^ population within the subset of CD86^+^ cDCs. This enabled us to use CD86^hi^MHCII^hi^ (Figure 5B) as the designation for activated and mature DCs. Using this designation, we determined that BH stimulates an increase in the number of activated CD86^hi^ cDC1s (Figure 5D) and cDC2s (Figure 5G), as well as a greater proportion of activated cDC1s (Figure 5E) and cDC2s (Figure 5H). Overall, these results demonstrate that BH enhances the activation of both cDC subsets in TDLN.

**Figure 5.**
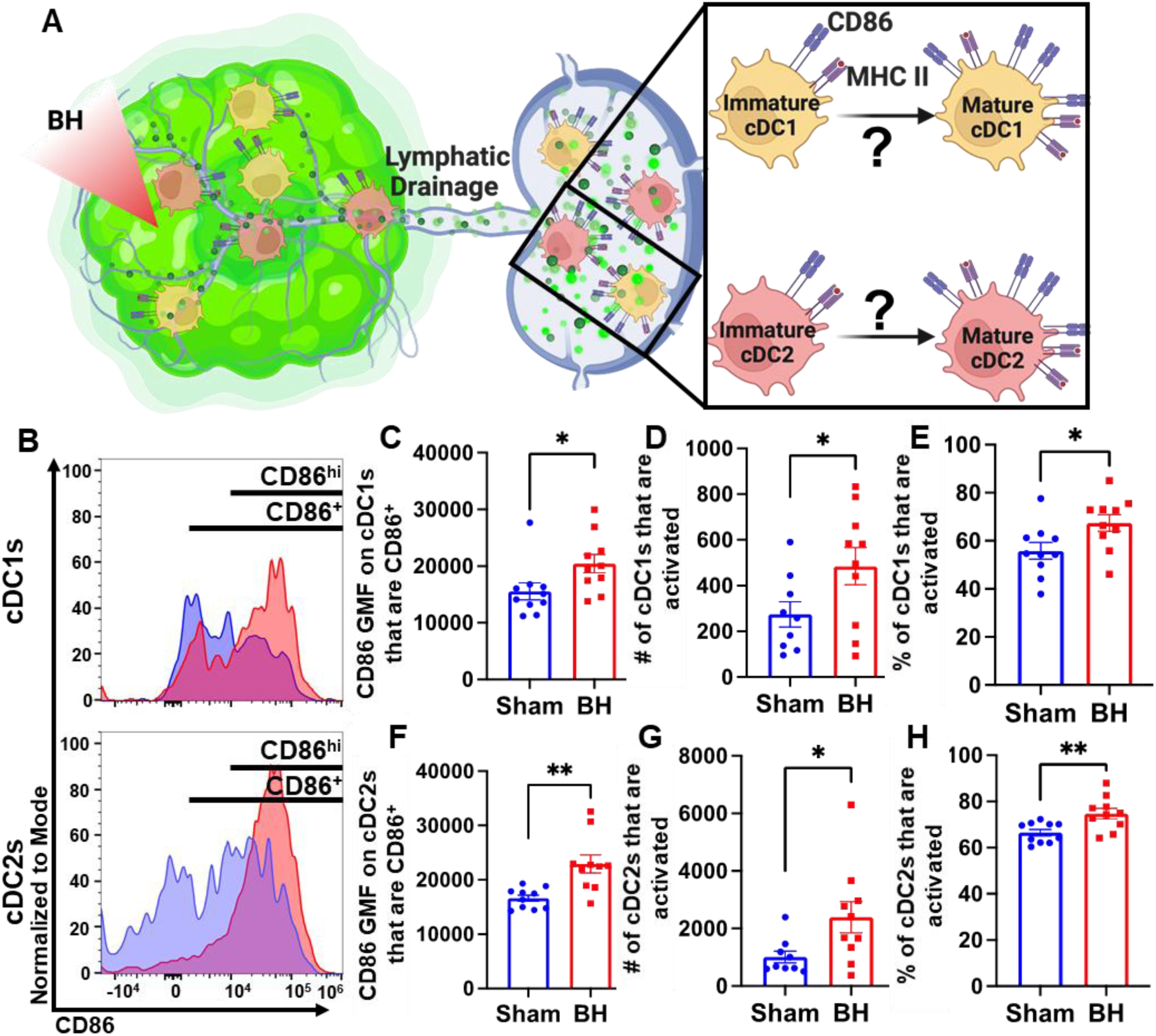
BH activates cDCs in tumor draining lymph nodes. **A.** Diagram illustrating the question of whether BH activates cDC1s and cDC2s. **B.** Frequency plots of side scatter vs CD86 for cDC1s (top) and cDC2s (bottom). **C & F.** Bar graphs of CD86 GMF of CD86^+^ cDC subsets. **D & G.** Bar graphs of number of activated cDCs (CD86^hi^MHCII^hi^). **E & H.** Bar graphs of percent of cDCs that are activated (CD86^hi^MHCII^hi^). n=10 per group. Unpaired Welch’s t-test: ∗ p < 0.05, ∗∗ p < 0.01. Means ± SEM.

### Conventional Dendritic Cell Activation by Boiling Histotripsy Depends on Tumor Antigen Acquisition

Because we observed that BH elicits increased total DC antigen acquisition and cDC activation, we asked whether BH-induced cDC activation was dependent on ZsG tumor antigen acquisition (Figure 6A). If not, and ZsG^-^ cDC also exhibit increased activation with BH, it would suggest that BH treatment liberates immunostimulatory molecules that are available to all cDC. To address this question, we analyzed CD86 expression on ZsG^-^ cDC subsets (Figure 6B). We observed no differences in cDC CD86 expression (Figure 6C and F) or changes in the number (Figure 6D and G) and percentages of activated ZsG^-^ cDCs (Figure 6E and H) between sham control and BH treated cohorts. These results indicate that BH alone is not inducing the cDC activation that we observed in Figure 5. Instead, activation only occurs as a consequence of ZsG acquisition.

**Figure 6.**
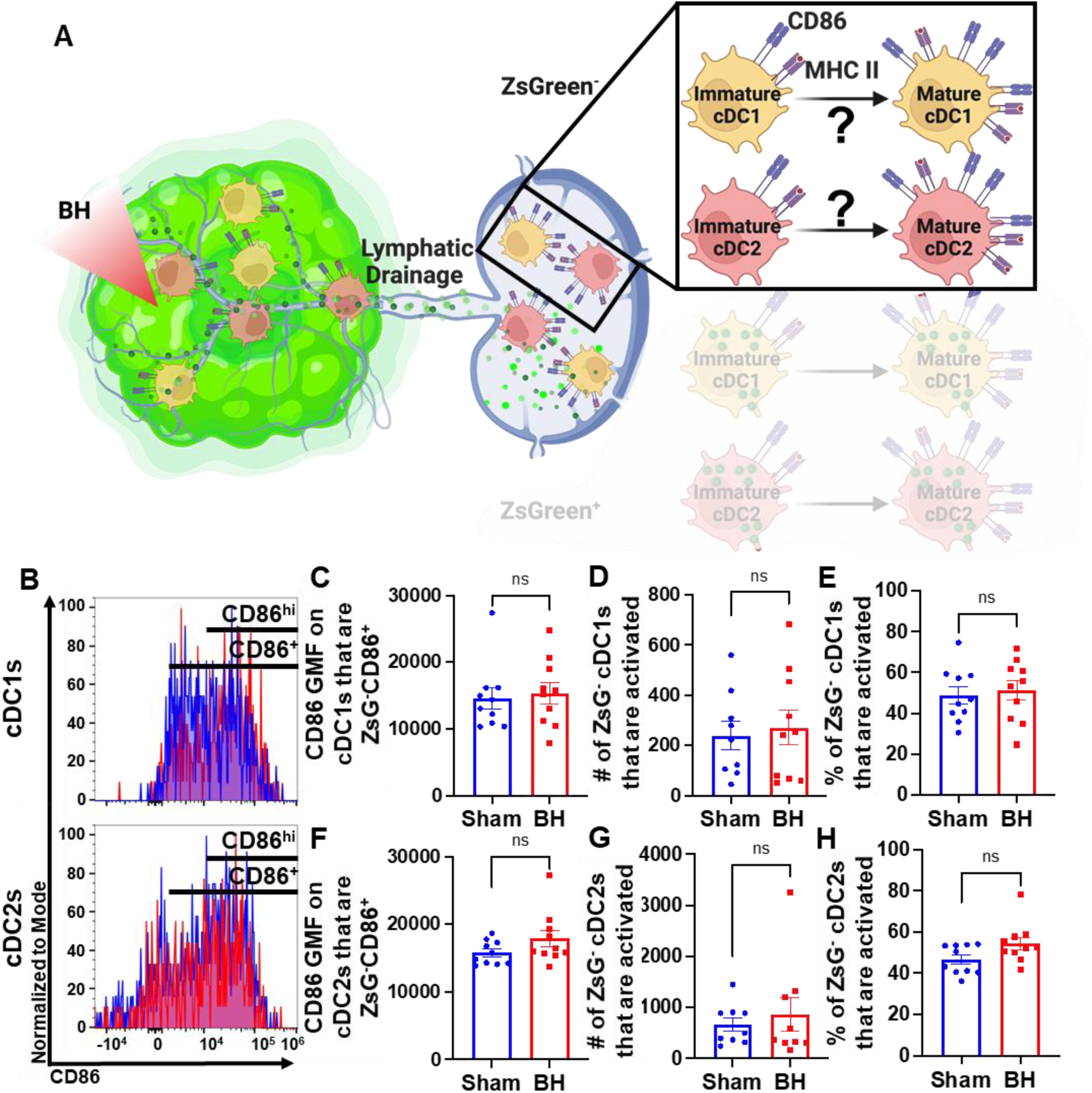
BH-induced cDC depends on ZsG acquisition. **A.** Diagram illustrating the overall question of whether ZsG acquisition is required for cDC1 and/or cDC2 activation. Data in this figure interrogate ZsG^-^ cDCs, so the ZsG^+^ portion is shaded. **B.** Frequency plots of side scatter vs CD86 for ZsG^-^ cDC1s (top) and ZsG^-^ cDC2s (bottom). **C & F.** Bar graphs of geometric mean fluorescence (GMF) intensity of CD86 on CD86^+^ZsG^-^ cDC subsets **D & G.** Bar graphs of number of specified ZsG^-^ cDC subset that are activated (CD86^hi^MHCII^hi^). **E & H.** Bar graphs of percent of specified cDC subset that are activated (CD86^hi^MHCII^hi^). n=10 per group. Unpaired Welch’s t-test: not significant. Means ± SEM.

### Tumor Antigen Acquisition Promotes Conventional Dendritic Cell Activation

We next interrogated ZsG^+^ cDC1s and cDC2s to understand the extent to which antigen acquisition was responsible for stimulating cDC activation (Figure 7A). While the majority of ZsG^+^ cDC1s and cDC2s are activated at baseline (Figure 7B and 7C), BH did significantly increase the percentage of ZsG^+^ cDC2s that are activated (Figure 7C). Next, we compared ZsG^-^ and ZsG^+^ cDC subsets to ascertain whether ZsG alone is capable of eliciting cDC activation. Importantly, we compared these populations in the same TDLN to account for any potential differences in response to BH. The baseline presence of ZsG, in the absence of BH, correlates with higher CD86 expression on cDC1s (Figure 7D) and cDC2s (Figure 7H). However, BH further increased CD86 levels on ZsG^+^ cDC1s (Figure 7E) and ZsG^+^ cDC2s (Figure 7I). This suggests there is a qualitative difference in cDC activation after tumor antigen acquisition as a consequence of BH.

**Figure 7.**
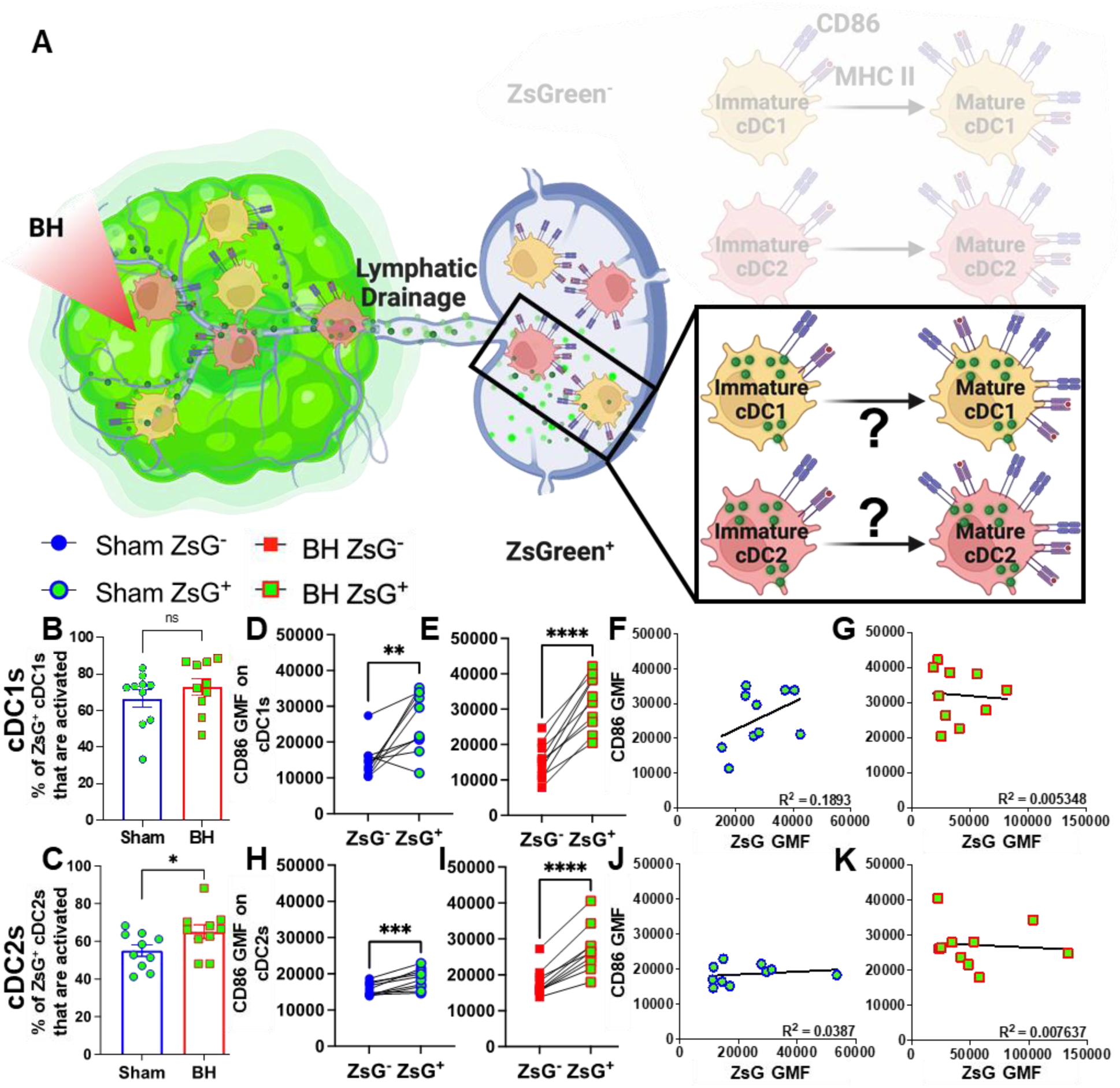
ZsG antigen promotes cDC maturation. **A.** Diagram illustrating the overall question of whether ZsG acquisition is required for cDC1 and/or cDC2 activation. Data in this figure interrogate ZsG^+^ cDCs, so the ZsG^-^ portion is shaded. **B & C.** Bar graphs of percent of ZsG^+^ cDC1s (**B**) and cDC2s (**C**) that are activated (CD86^hi^MHCII^hi^). n=10 per group. Unpaired Welch’s t-test: ∗ p < 0.05. **D-E & H-I.** Geometric mean fluorescent (GMF) intensity of CD86 on ZsG^-^ and ZsG^+^ cDC1s (**D-E**) and cDC2s (**H-I**). n = 10. Paired t-test: ∗ p < 0.05, ∗∗ p < 0.01, ∗∗∗ p < 0.001, ∗∗∗∗ p < 0.0001. **F & J.** CD86 expression as a function of ZsG expression in ZsG^+^ cDC1s (**F**) and ZsG^+^ cDC2s (**J**) after sham treatment. **G & K.** CD86 expression as a function of ZsG expression in ZsG^+^ cDC1s (**G**) and ZsG^+^ cDC2s (**K**) after BH. Correlations are by linear regression with displayed R^2^ values.

To better understand the quality of BH-induced cDC activation, we compared BH-induced CD86 expression levels in cDCs to those elicited by administration of a TLR3 agonist (i.e., polyinosinic-polycytidylic acid with poly-L-lysine double-stranded RNA [polyI:CLC]) that is known to be a highly potent driver of cDC activation (Figure S8). In this experiment, ZsG^+^ cDCs in TDLNs of saline-treated control mice also exhibited elevated CD86 expression when compared to ZsG^-^ cDCs (Figure S8A and S8C). From there, as expected, PolyI:CLC massively increased CD86 expression on the surface of cDCs. Furthermore, the trend that ZsG^+^ cDCs express higher levels of CD86 in response to BH was maintained with polyI:CLC treatment (Figure S8B and S8D). Thus, while BH elicits cDC activation in an antigen-dependent manner, the magnitude of the activation response does not match that generated with direct TLR3 agonism.

We then examined whether the increased activation that accompanies ZsG presence in cDCs after BH is reflective of the amount of ZsG acquisition or whether there is a qualitative difference in ZsG with respect to cDC activation. We hypothesized that if increases in the activation of ZsG^+^ cDCs observed after BH were simply a function of acquiring more antigen, a greater amount of acquired ZsG (i.e., ZsG GMF) would lead to greater expression levels of CD86. Nonetheless, we found no correlation between the amount of ZsG in either cDC1 or cDC2 and the level of CD86 expression for either sham (Figure 7F and J) or BH (Figure 7G and K) treated mice. Therefore, while the presence of ZsG tumor antigen presence correlates with CD86 expression on the surface of cDCs, the lack of correlation between the amount of ZsG acquired and the surface level expression of activation marker CD86 suggests that additional stimuli, such as a DAMP(s), are complexed with ZsG after BH and are required for the elevated CD86 expression.

### Boiling Histotripsy Does Not Alter Total DCs or Migratory Proportions of cDC1s

There is evidence that BH can augment DC migration to the TDLN [25], but it is not known whether cDCs acquire antigen intratumorally or in the TDLN (Figure 8A). We addressed this question two ways. First, we examined the total number of DCs in the TDLN (Figure 8B). We observed no change in DC representation (Figure 8C and 8D), which is consistent with a lack of cDC migration to TDLN. Second, we examined the representation of both tissue-resident (CD8α^+^) and migratory (CD103^+^) cDC1s (Figure 8E) in TDLN. Here, we observed no changes in (i) the numbers of CD8α^+^ and CD103^+^ cDC1s (Figure 8F and 8G), (ii) percentages of ZsG^+^ cDC1s that are CD8α^+^ and CD103^+^ (Figure 8J and 8K), and (iii) percentage of cDC1s that are CD103+ (Figure 8I). These findings are again consistent with a lack of cDC migration to TDLN in response to BH. Though the percentage of tissue-resident cDC1s decreased modestly with BH (Figure 8H), we do not think this is biologically significant as the migratory cDC1 proportion did not change with BH (Figure 8I). Additionally, both tissue-resident (Figure 8L) and migratory (Figure 8M) cDC1s exhibited an increase in the proportion of cells that acquired ZsG^+^ significantly. In all, these results suggest that the increase in ZsG-tumor antigen observed for cDC1s is due to the acquisition of cell-free tumor antigen that flows to the TDLN after being liberated by BH.

**Figure 8.**
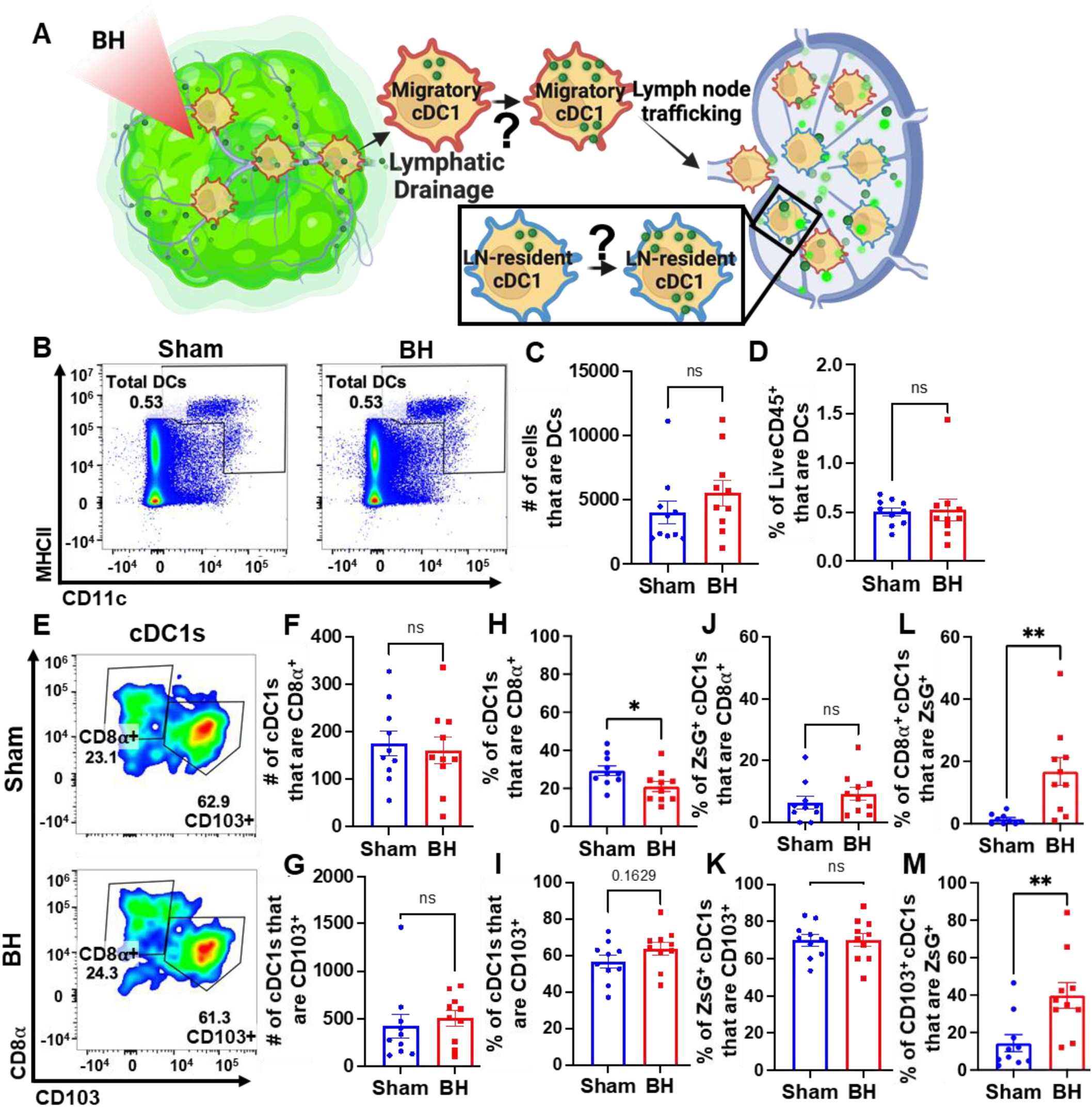
BH changes neither total DCs nor the proportions of ZsG^+^ cDC1s that are LN-resident (CD8α^+^) and migratory (CD103^+^). **A.** Diagram illustrating the question of whether cDC1s acquire BH-liberated ZsG in tumor and/or TDLN. **B.** Density scatter plots of CD11c vs MHCII with DCs (CD11c^+^MHCII^+^) in sham and BH treated TDLNs. **C.** Number of total DCs. **D.** Percentage of LiveCD45^+^ cells that are DCs. **E.** Density scatter plots of CD8α vs. CD103 to identify tissue-resident (CD8α^+^CD103^-^) and migratory (CD103^+^) cDC1s in TDLNs of sham and BH treated mice. **F & G**. Number of tissue-resident (**F**) and migratory (**G**) cDC1s. **H & I.** Percentage of cDC1s that are tissue-resident (**H**) and migratory (**I**). **J & K.** Percentage of ZsG^+^ cDC1s that are tissue-resident (**J**) and migratory (**K**). **L & M.** Proportion of tissue-resident (**L**) and migratory (**M**) cDC1s that are ZsG^+^. n=10. Unpaired Welch’s t-test: ∗ p < 0.05, ∗∗ p < 0.01. Means ± SEM.

## Discussion

The intent of this study was to fill crucial gaps in our understanding of how BH affects key elements of the cancer immunity cycle, in particular the relationship between tumor ablation and tumor antigen acquisition by cDCs, and the allied activation of cDCs, both of which are critical to the subsequent activation of tumor-specific T cells. By deploying a sparse scan BH treatment regimen that yields abscopal control of B16F10 melanoma tumors expressing a ZsG model antigen, we were able to make the first ever direct measurements of (i) the dynamics of tumor antigen trafficking to TDLNs after BH, (ii) the identity of immune cell types that acquire BH-liberated antigen, including antigen partitioning amongst cDCs, (iii) how cDC maturation is affected by BH and the role of antigen acquisition in this process, and (iv) whether tumor antigen is dominantly acquired by cDCs in the tumor or TDLN. We observed a striking increase in tumor antigen presence in CD45^+^ immune cells in the TDLN 24h after BH, which abates by 96h post-ablation. Within TDLNs, B cells, macrophages, monocytes, and granulocytes, as well as both cDC1s and cDC2s, all acquired markedly more tumor antigen after BH. Notably, BH drove significant activation of both cDC subsets, with a more detailed analysis of our flow cytometry data revealing that (i) cDC activation was dependent upon tumor antigen acquisition and (ii) the tumor antigen liberated by BH is likely complexed with a DAMP(s). Because the increase in tumor antigen-bearing cDCs in TDLN after BH did not correlate with an increase in total DC presence in the TDLN or a marker of cDC migration (CD103), we posit that cDCs do not traffic to the TDLN with antigen, but rather acquire antigen as it flows through afferent lymph vessels into the TDLN. In all, our results illuminate numerous previously unknown features of how BH, applied with a monotherapy protocol that elicits abscopal tumor control, drives tumor antigen trafficking to TDLN and instructs DC function. Going forward, such information will be invaluable for rationally tuning BH treatments for optimal immunological tumor control and selecting immunotherapies, as well as their administration timings, for improved combination treatments.

### Dynamics of Tumor Antigen Trafficking to the TDLN

We determined that BH drives a nearly 3-fold increase in the proportion of antigen positive immune cells at 24 h, with a return to baseline by 96 h after BH treatment. We chose 24 h as the timepoint in all subsequent studies based on this finding. We also emphasize that this time point is commonly used in other studies of antigen and DC trafficking [26–29] and is appropriate for this particular application. Indeed, the choice of the 24h timepoint permits identification of both small soluble antigens, such as ZsG (26 kDa m.w.) that reach the TDLN in an acellular fashion within minutes [26–28,30,31], as well as DCs that acquire antigen in the tumor microenvironment and may take ∼18 hours to traffic to the TDLN [32]. Notably, molecules exceeding ∼60-70 kDa appear to need a cell (e.g., migratory CD103^+^ cDC1s) to traffic the antigen from peripheral tissues (e.g., tumor) to the TDLN [27,28], thus our use of the relatively small ZsG antigen permits assessment of antigen trafficking via both means.

We also provide considerable evidence that tumor antigen acquisition after BH is independent of cDC trafficking from the BH-treated tumor to the TDLN. First, enhanced tumor antigen presence was observed after BH in cell types that do not migrate from the tumor to TDLN (e.g., B cells). Second, within the cDC1 and cDC2 subsets, we observed no changes in total cDCs, nor in cDCs containing ZsG. This result suggests that cDCs are acquiring tumor antigen in the TDLN. Third, when we quantified CD8α^+^ (tissue-resident) and CD103^+^ (migratory) cDC1s, we found neither an increase in the CD103^+^ migratory population in TDLN in response to BH, nor an increase in the presence of ZsG tumor antigen in these cells. Together, these data strongly argue that BH treatment does not promote the migration of cDC to TDLN. Rather, cell-free tumor debris is reaching the TDLN. That said, these results do run counter to another study wherein BH increased the numbers of total and transferred (i.e., injected intratumoral CSFE-labelled bone-marrow derived DCs [BMDCs] two days post BH treatment) DCs in the TDLN in the context of MC-38 colorectal cancer [25]. The most obvious difference between our studies is the difference in tumor model (i.e., 3-4×10^5^ B16F10-ZsG vs 1×10^6^ MC-38 [25]). At baseline, MC-38 grows slower and has higher T cell, NK cell and cDC infiltration. Moreover, MC-38 tumors have a superior response to anti-PD-1 therapy [33]. A more nuanced difference appears in the number of BH application points per tumor. While our treatments entailed 20 to 79 sonications per tumor, Hu et al. applied 12 to 16 sonications [25]. Our BH ablation regimen appears to be more aggressive given similarities in focal size and transducer frequency. These results may indicate that DC-sparing ablation regimens can be crafted to better promote DC trafficking to the TDLN, though the therapeutic necessity of such DCs has yet to be determined. An alternate hypothesis is that intratumoral DCs do not play a significant role in the immunological response to BH in melanoma. In that case, sparing DCs from BH ablation will confer no benefit. Thus, increasing the intensity and/or fraction of BH ablation to liberate more tumor antigen may further augment favorable responses. Another caveat to this interpretation is that we only examined cDCs 24h after BH. It is possible that a small number of migratory cDCs emerge later, although we determined that no increase in tumor antigen in TDLN is evident at 96 post BH. These results have important implications for choosing tumor models, BH ablation fractions, and timepoints in future studies aimed at combining BH with immunotherapies.

### Partitioning of Tumor Antigen in cDCs in TDLN

Another important objective of our studies was to determine whether BH-liberated tumor antigen is preferentially acquired by either cDC1s or cDC2s in the TDLN, as this may influence the subsets of T cells primarily stimulated by BH. We found an increase in the number and proportion of antigen positive cells in both cDCs. Because cDC1s primarily activate CD8^+^ T cells [17–19], while cDC2s are required for initial priming of CD4 T cells [17], these results indicate that we should not expect a biasing toward CD4^+^ or CD8^+^ T cell activation due to uneven cDC tumor antigen acquisition. Interestingly, the cDC2 subset did exhibit more antigen per cell, which may suggest that cDC2s express phagocytosis receptors that are more adept at acquiring BH-liberated antigen and/or that cDC2s are preferentially positioned in the TDLN (i.e., close to the lymphatic cannulae) to acquire this antigen.

### Activation of cDC in TDLN as a Function of BH and Tumor Antigen Acquisition

Our studies have revealed unexpected relationships between BH-mediated tumor antigen liberation and the activation state of cDCs in the TDLN. Indeed, we found that only ZsG^+^ cDCs exhibit increased CD86 expression as a function of BH treatment. Given that previous studies have documented the release of DAMPs capable of driving CD86 expression on BMDC in vitro [34], as well as antigen-agnostic activation of DCs after BH in mouse lymphoma [12], we had expected a similar global activation of cDCs independent from the acquisition of tumor antigen. The current data suggests that either the process of acquiring tumor antigen drives cDC activation or that stimulatory molecules complexed with the tumor antigens are responsible for cDC activation. Moreover, we determined that the level of CD86 expression was higher on ZsG^+^ cDC from BH treated TDLN compared to sham controls, yet we did not observe a proportional increase in CD86 expression as cDC acquired more tumor antigen after BH. The tentative conclusion from these observations is that the increased level of cDC activation seen after BH is not simply a function of there being more tumor antigen available to engulf. Instead, exposure to BH may modify the tumor antigen in a manner that promotes cDC activation, perhaps by complexing it with a DAMP.

### Broader Implications for Boiling Histotripsy-Driven Antigen Trafficking

To the best of our knowledge, our study is the first to track tumor antigen in the TDLN after its liberation from a solid tumor by BH. It is reasonable to hypothesize that, in studies by other investigators wherein BH elicited DC activation and migration [11,12,16] and/or enhanced T cell representation in solid tumors [11,13,15,16], similar tumor antigen trafficking to TDLN occurred. Yet, the extent to which such trafficking may occur is likely dependent upon several factors. These include (i) BH ablation spacing and fraction, (ii) mechanical properties of the solid tumor, and (iii) the quality of lymphatic drainage from the solid tumor to the TDLN. Comparisons of tumor stiffness and lymphatic quality are difficult to make between published studies; however, BH parameters are accessible. Here, we used a 1 mm BH ablation spacing (Figure S1). Other studies share, to some extent, this general characteristic. For example, 1 mm BH treatment spacing has been used to generate immunological responses consistent with antigen trafficking to TDLN in immunogenic MC-38 colon adenocarcinoma [11] and EG.7-OVA lymphomas [12], with 1-2 mm spacing showing efficacy in E0771 and MM3MG-HER2 breast tumors [13]. That said, if we instead consider BH ablation fraction (20% in our study; Figure S1), a wider range of effective values has been reported. Indeed, BH ablation fractions for studies showing augmented DC activation and/or T cell representation range from 2% for neuroblastoma [14], to 20%-40% for breast tumors [13] and 40%-50% for B16F10 melanoma [15]. Moreover, there is also evidence that the intensity of BH treatment within a single focal spot may be important [16]. We submit this is a factor which could be become more significant when treating dense stromal tumors, such as 4T1 breast tumors. In contrast, relatively soft B16F10-ZsG tumors were studied here. When this discussion is considered in light of our data suggesting that increasing ablation fraction could be beneficial (Figure 8), we submit that tuning BH parameters for optimal tumor antigen release from different solid tumor types is an important topic of future investigation for this field.

## Materials and Methods

### Cell line and animal maintenance

The B16F10-ZsGreen cell line was a kind gift from Dr. Matthew Krummel at the University of California, San Francisco [35]. Cells were maintained in RPMI-1640+L-Glutamine (Gibco #11875-093) supplemented with 10% Fetal Bovine Serum (FBS, Gibco #16000-044) at 37°C and 5% CO_2_ (Thermo Fisher Scientific, Heracell 150i Cat#51-032-871). Thawed cells were cultured for up to three passages and maintained in logarithmic growth phase for all experiments. Cells tested negative for mycoplasma prior to freezing.

All mouse experiments were conducted in accordance with the guidelines and regulations of the University of Virginia and approved by the University of Virginia Animal Care and Use Committee. Eight-week-old to ten-week-old male C57Bl/6J mice were obtained from The Jackson Laboratory (Jax #000664). 3-4×10^5^ B16F10-ZsGreen cells were implanted subcutaneously (s.c.) into the right flank of mice after shaving through a 25G x 1 ½ in needle (BD PrecisionGlide Needle #305127). For the growth control and survival study, 4×10^5^ B16F10-ZsGreen cells were s.c. implanted into the right and left flanks of mice and treated with sham/BH 14 days post-inoculation. Mice were housed on a 12- hour/12- hour light/dark cycle and supplied food ad libitum. Tumor outgrowth was monitored via digital caliper measurements. Tumor volume was calculated as follows: volume = (length×width^2^)/2. Thirteen- or fourteen-days following tumor implantation, mice were randomized into groups in a manner that ensured matching of mean starting tumor volume across experimental groups.

### In vivo ultrasound-guided boiling histotripsy

Mice underwent sham or BH treatment 13- or 14-days post-inoculation. On treatment day, mice were anesthetized with an intraperitoneal (i.p.) injection of ketamine (50 mg/kg; Zoetis) and dexdomitor (0.25 mg/ kg; Pfizer) in sterilized 0.9% saline (Hospira #PAA128035). Dexdomitor was reversed with a s.c. injection of atipamezole hydrochloride (0.25 mL in 10 mL saline, 0.4 mL s.c., Antisedan, Zoetis) after sham or BH treatment. Right flanks of mice were shaved, after which BH was performed using an in-house built ultrasound-guided FUS system. This includes incorporation of ultrasound visualization/guidance orthogonal to the focal axis of the therapy transducer. The system uses one of two linear imaging arrays: 1) Acuson Sequoia 512, 15L8 imaging probe, 8 MHz, 25 mm field (Siemens, Inc.) width or 2) Acuson S2000 Helix Evolution Touch, 14L5 SP imaging probe, 10 MHz, 25 mm field width (Siemens, Inc.). A 1.1 MHz center-frequency, single-element therapy transducer H-101 (Sonic Concepts Inc., Bothel, WA) was used in combination with an arbitrary function generator (Tektronix, AFG 3052C) and amplifier (E&I, 1040L) to produce BH treatments. This therapy transducer had an active diameter of 64 mm and radius of curvature of 63.2 mm (i.e., the geometric focal distance). The transducer was operated at third harmonic (3.28 MHz), with a -6dB focal size of 0.46 mm x 0.46 mm x 3.52 mm = ∼0.39 mm^3^. Both the imaging and treatment transducers were ultrasonically coupled to the animal using degassed, deionized water at 37°C during the duration of each BH treatment. BH was applied in a pulsed fashion for 10 s, at a peak negative pressure = 21 MPa, pulse repetition frequency = 4 Hz, pulse length = 3 ms, with treatment points spaced 1 mm in a rectangular grid pattern and 2 planes of treatment, which were separated by 2 mm. With this ablation pattern and focal size, we calculate that ∼20% of each tumor was exposed to BH. The treatment scheme is outlined in Figure S1. Sham treatment comprised of fully submerging the flank tumor in the 37°C water bath for 6 minutes.

### PolyI:CLC delivery

PolyI:CLC (Oncovir, Inc., Hiltonol®) was injected i.p. at 13 days post-inoculation with 75 μg/0.1 mL diluted with sterilized 0.9% saline. Flow cytometry was performed 24 hr after injection.

### Flow Cytometry

At 13 days post-tumor inoculation, tumor draining lymph nodes (TDLNs) - axial and brachial on the right side – as well as contralateral non-tumor draining lymph nodes were excised and pooled. LNs were subjected to manual homogenization (Wheaton, Tenbroeck Tissue Grinder #62400-518) and filtered through 100 μm filter mesh (Genesee Scientific # 57-103) to generate single-cell suspensions, which were then washed in 1X PBS, centrifuged at 1200 RPM for 5 minutes (Eppendorf 5180) and stained for cell viability using Fixable Live/Dead Blue for 30 min at 4°C. Next, the samples were exposed to anti-mouse CD16/32 to block Fc gamma receptors for 15 min at 4°C. Afterwards, cells were washed with FACS buffer, centrifuged, and resuspended in a mixture of Brilliant Stain Buffer and FACS+2% normal mouse serum (Valley Biomedical, Inc., #AS3054) at a ratio of 1:9, respectively, and stained for 30 min at 4°C with fluorescent monoclonal antibodies for CD45, CD11b, Ly-6G, Ly-6C, F4/80, CD11c, MHCII, XCR1, SIRPα, CD19, CD3, CD8α, CD86, CCR7 and CD103. Antibody clone information, supplier name and catalog number can be found in Table S2. Lastly, cells were fixed in 1X BD FACS Lysis for 10 min at room temperature, and then resuspended in FACS buffer for running. Flow cytometry was performed with the Cytek Aurora Borealis (Cytek Biosciences) and SpectroFlo v3.0.3 software (Cytek Biosciences). Data was analyzed using FlowJo 10 software (FlowJo, LLC). All gating strategies can be found in Figures S2, S4, S6 and S8.

### Statistical Analyses

Most statistical analyses were performed in GraphPad Prism 9 (GraphPad Software). Mouse survival was analyzed using a Kaplan-Meier analysis and a log-rank (Mantel-Cox) test was used to assess significance. When comparing two groups of flow cytometry data or area under the curve (AUC; i.e., sham vs BH), an unpaired, two-tailed t-test with Welch’s correction (i.e., did not assume equal standard deviations) was performed. A paired t-test was used to compare within group differences based on ZsG positivity (i.e., sham ZsG^-^ vs sham ZsG^+^; BH ZsG^-^ vs BH ZsG^+^). Groups of flow cytometry summary data across time were compared using a full-model, two-way analysis of variance (ANOVA) and Tukey post-hoc tests to assess significance of factors (i.e., time [factor 1] and sham/BH [factor 2]) and between individual groups, respectively. All figures show the mean ± standard error of the mean (SEM). P-values and significance are specified in figure legends. All figure schematics were made with BioRender.com.

Tumor growth data was modeled in MATLAB 2022b using non-linear least squares with a logistics model [36–38] out to day 35 post-inoculation to account for mouse drop out (i.e., tumor size met one of the humane endpoint criteria) for each individual mouse. The resulting curves were averaged together. The modeled data was appended to actual tumor data up until day 35 (e.g., if the mouse dropped out at day 25, only days 26 through 35 of the modeled data are used). The following variation of a logistic model was used (Eq. 1).

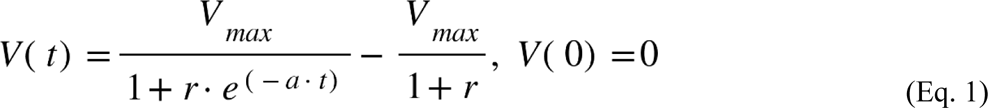

The fitted parameters are “*r*” and “*a*” while, if the maximum tumor volume of the raw data is less than 4000 mm^3^, *V_max_* is 4000 mm^3^, otherwise, *V_max_* is set to the maximum tumor volume. The value of 4000 mm^3^ was chosen because this is roughly the largest volume a tumor can achieve given the humane endpoint criteria. The R^2^, “*r*” and “*a*” values can be found in Table S1. Comparisons of these logistic average tumor curves between treatment groups were performed with a full model, two-way repeated-measures ANOVA with two factors (i.e., time [repeated-measures] and sham/BH) and corresponding interaction terms using the Geisser-Greenhouse correction. The fixed effect of BH treatment was used to determine significance. Furthermore, to summarize the logistic average tumor growth curves with a single parameter, we calculated the AUC from day 14 to 35 for the averaged curves using the trapezoid rule [39].

## Acknowledgments

We thank the Flow Cytometry Core Facility of the University of Virginia for the use of their cytometers. This core is supported by the National Cancer Institute P30-CA044579 Center Grant. We thank Dr. Matthew Krummel for his gift of the B16F10-ZsGreen cell line.

## Author contributions

EAT and LEK contributed equally to all areas.

Conceptualization: EAT, LEK, TNJB, RJP

Methodology: EAT, LEK, FRE, ASM, MRE, TNJB, RJP

Formal analysis and investigation: EAT, LEK, FRE, ASM

Writing - original draft preparation: EAT, LEK

Writing - review and editing: EAT, LEK, FRE, ASM, MRE, TNJB, RJP

Funding acquisition: TNJB, RJP

Resources: MRE, TNJB, RJP

Supervision: MRE, TNJB, RJP

## Data and materials availability

All data are available in the main text or supplementary materials and are available upon request.

