## Supplemental Information for "Focused ultrasound ablation of melanoma with boiling histotripsy yields abscopal tumor control and antigen-dependent dendritic cell activation"

Eric A Thim *et al.*

**This PDF file includes:**

Figs. S1 to S8
Tables S1 to S2

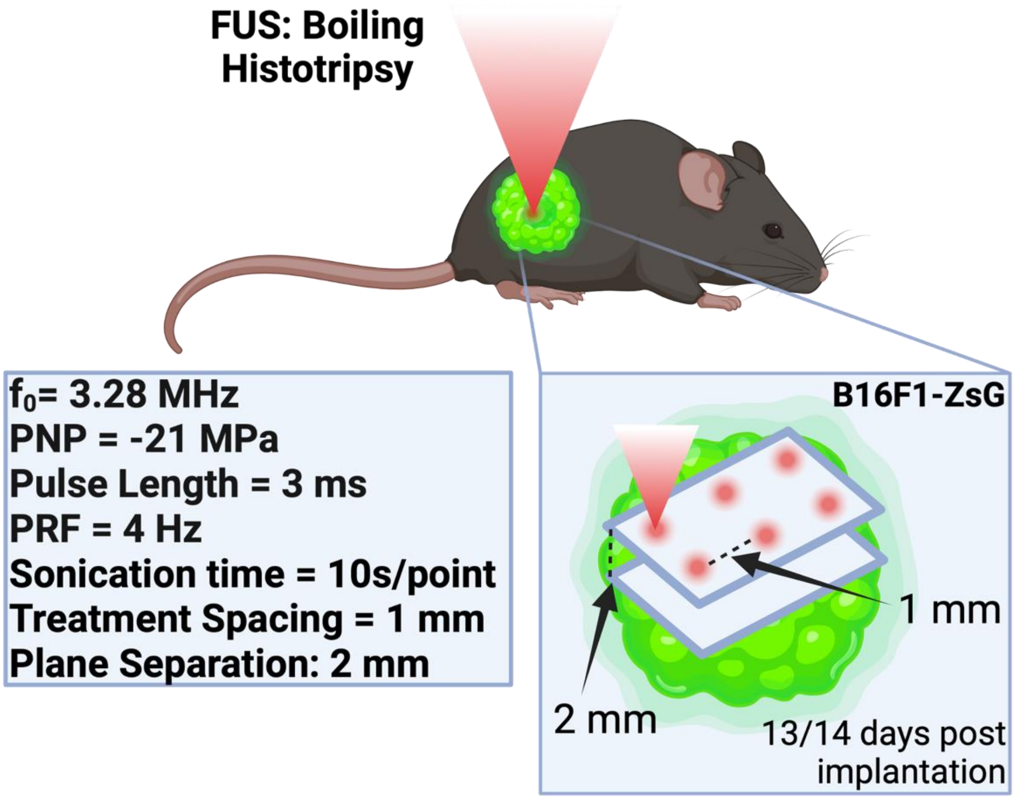


**Figure S1. Schematic for boiling histotripsy treatment.** Boiling histotripsy was applied with the following parameters: Operational frequency ($f_0$ = 3.28 MHz); peak-negative pressure (PNP = -21 MPa); pulse length (3 ms); pulse repetition frequency (PRF = 4 Hz); sonication time = 10 s/point; treatment spacing = 1 mm; plane separation = 2 mm. With this treatment scheme, ~20% of the tumor volume is within the -6dB focus during BH application.

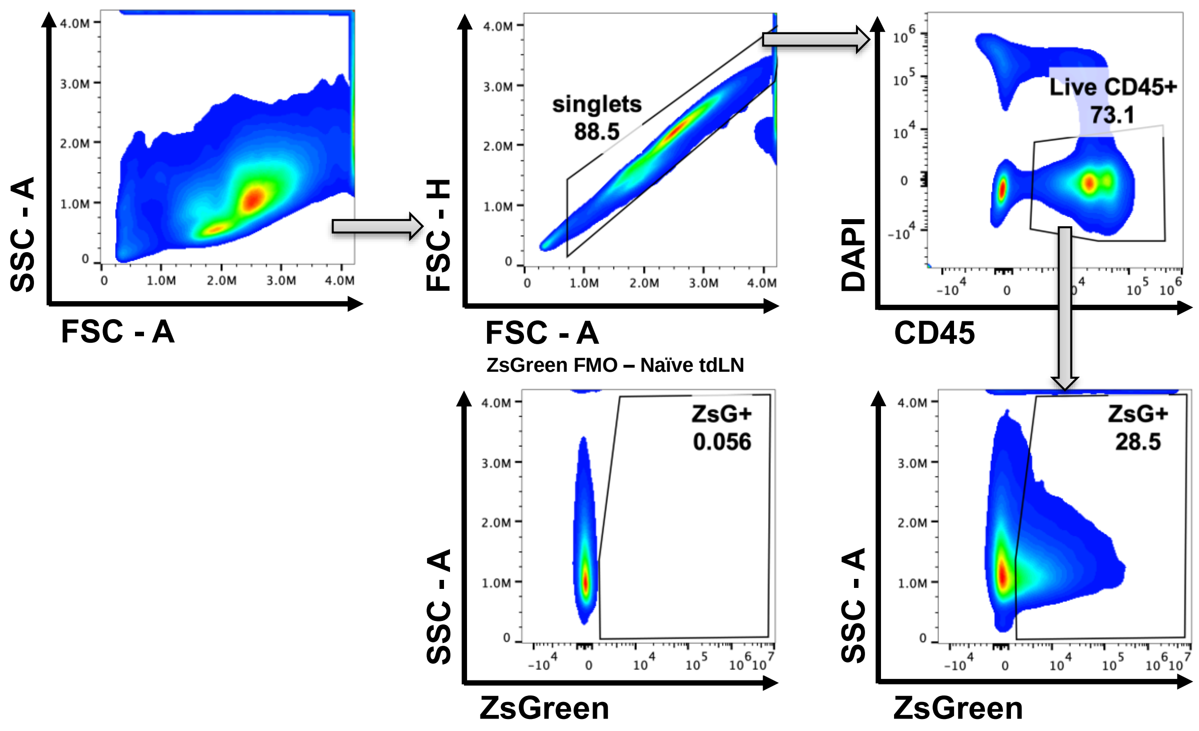


**Figure S2. Gating strategy for flow cytometry analysis of Figure 1.** ZsG-naïve mice were used as the gating control for ZsG positivity in LiveCD45$^+$ immune cells.

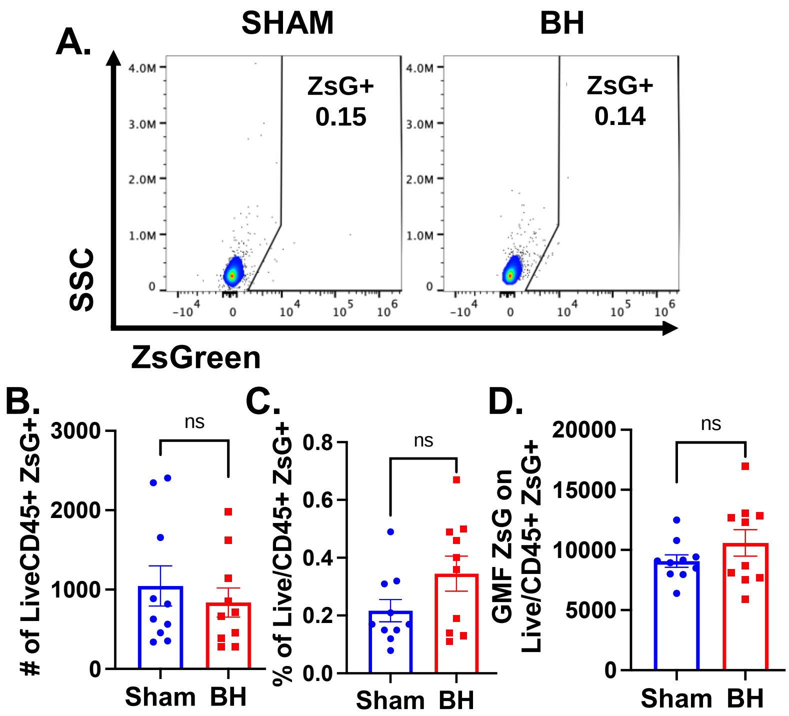


**Figure S3. BH does not promote ZsG tumor antigen acquisition in non-tumor draining lymph nodes 24 hrs post treatment. A.** Density scatter plots of side scatter vs ZsG indicating percent of Live/CD45$^{+}$ cells that are ZsG$^{+}$. **B-D.** Summary of flow cytometry analysis. **B**) Number of LiveCD45$^{+}$ZsG$^{+}$ cells. **C**) Percent of LiveCD45$^{+}$ cells that are ZsG$^{+}$. **D**) Geometric mean fluorescent intensity (GMF) of ZsG on LiveCD45$^{+}$ZsG$^{+}$ cells. n=10. Unpaired Welch's t-test: not significant. Means ± SEM.

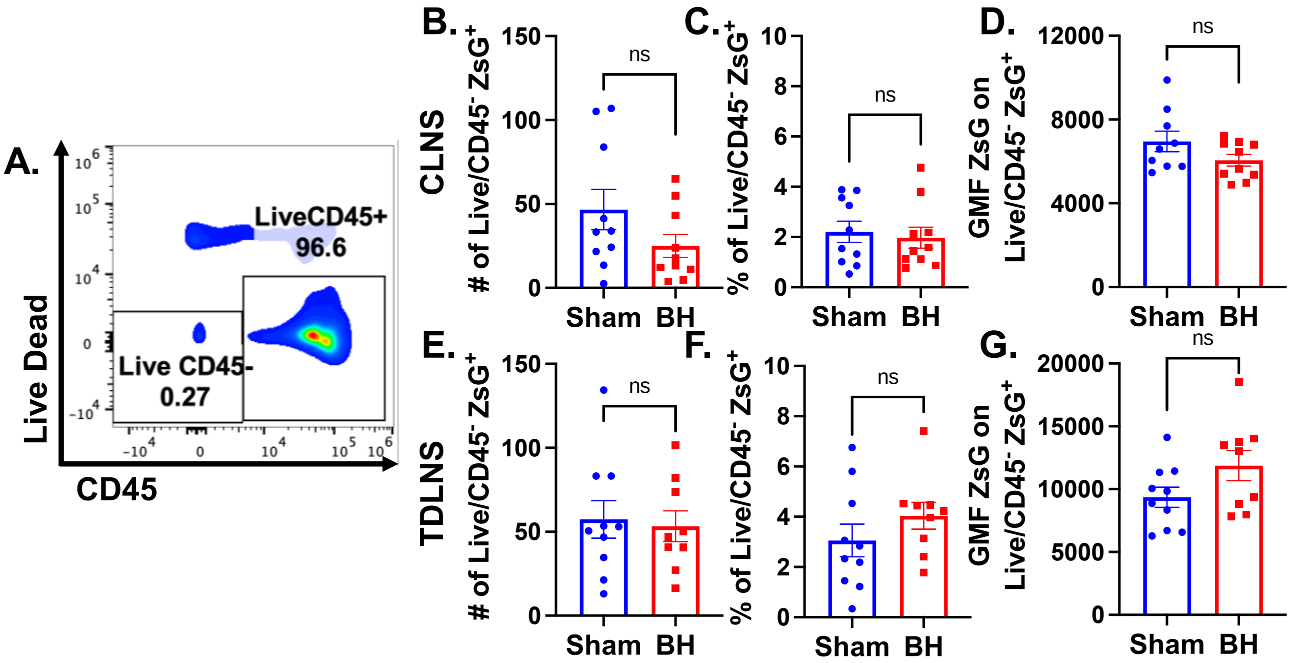


**Figure S4. BH does not promote metastatic spread of melanoma 24 hrs after treatment. A.** Density scatter plots of Live Dead vs CD45 indicating percent of singlet cells that are Live/$CD45^+$ and Live/$CD45^-$. The subsequent summary plots are in reference to $ZsG^+$ portion of $LiveCD45^-$ cells. **B, E)** Number of $LiveCD45^-ZsG^+$ cells in the contralateral and tumor draining lymph nodes. **C, F)** Percent of $LiveCD45^-$ cells that are $ZsG^+$ in CLNs and TDLN. **D, G)** The geometric mean fluorescent intensity (GMF) of ZsG on $LiveCD45^-ZsG^+$ cells in CLNs and TDLN**.** n=10. Unpaired Welch's t-test: not significant. Means ± SEM.

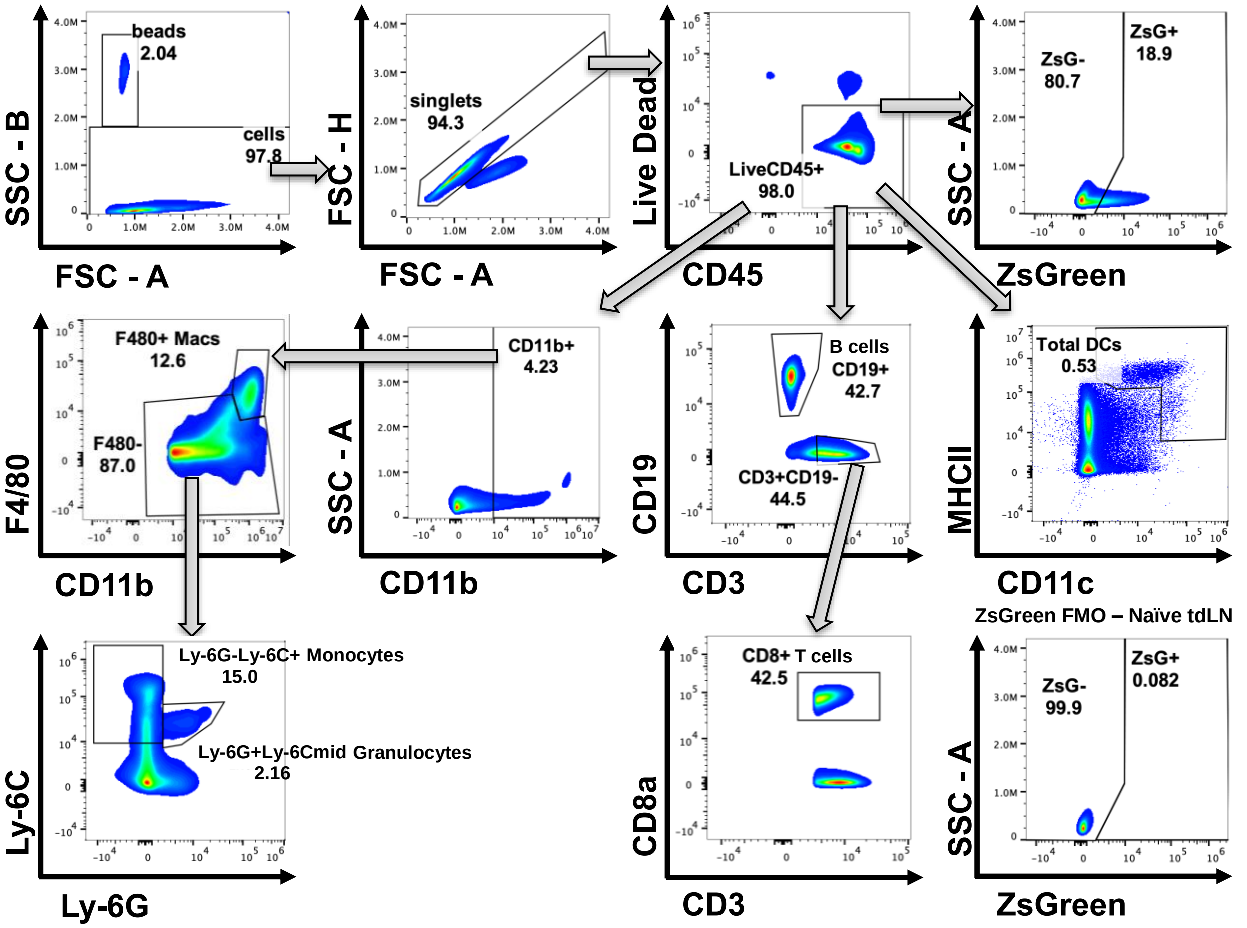


**Figure S5. Gating strategy for flow cytometry analysis of Figure 2 and Figure S3.** Gating strategy presented for the following immune cell subsets after cells and singlets: Macrophages (LiveCD45$^{+}$CD11b$^{+}$F4/80$^{+}$); Monocytes (LiveCD45$^{+}$CD11b$^{+}$F4/80$^{-}$Ly-6G$^{-}$Ly-6C$^{+}$); Granulocytes (LiveCD45$^{+}$CD11b$^{+}$F4/80$^{-}$Ly-6G$^{+}$Ly-6C$^{mid}$); B cells (LiveCD45$^{+}$CD19$^{+}$); CD8 T cells (LiveCD45$^{+}$CD3$^{+}$CD19$^{-}$CD8$^{+}$); Total DCs (LiveCD45$^{+}$CD11c$^{+}$MHCII$^{+}$). Gating strategy for ZsGreen$^{+}$ (ZsG$^{+}$) immune cells and the corresponding control from ZsG-naïve mice are outlined.

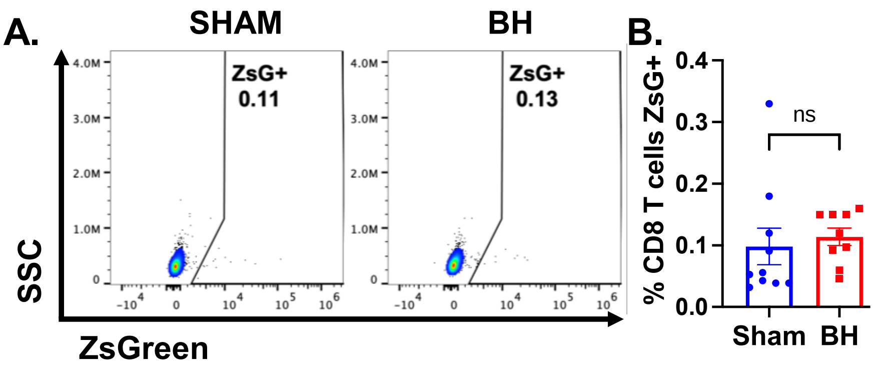


**Figure S6. Negative control for ZsG fluorescence showing antigen acquisition by non-APC and non-phagocytic cells (CD8$^+$ T cells). A.** Density scatter plots of side scatter vs ZsG indicating percent of LiveCD45$^+$CD19$^-$CD3$^+$CD8$^+$ T cells that are ZsG$^+$. **B.** Summary of flow cytometry analysis in **A**. n=10. Unpaired Welch's t-test: not significant. Means ± SEM.

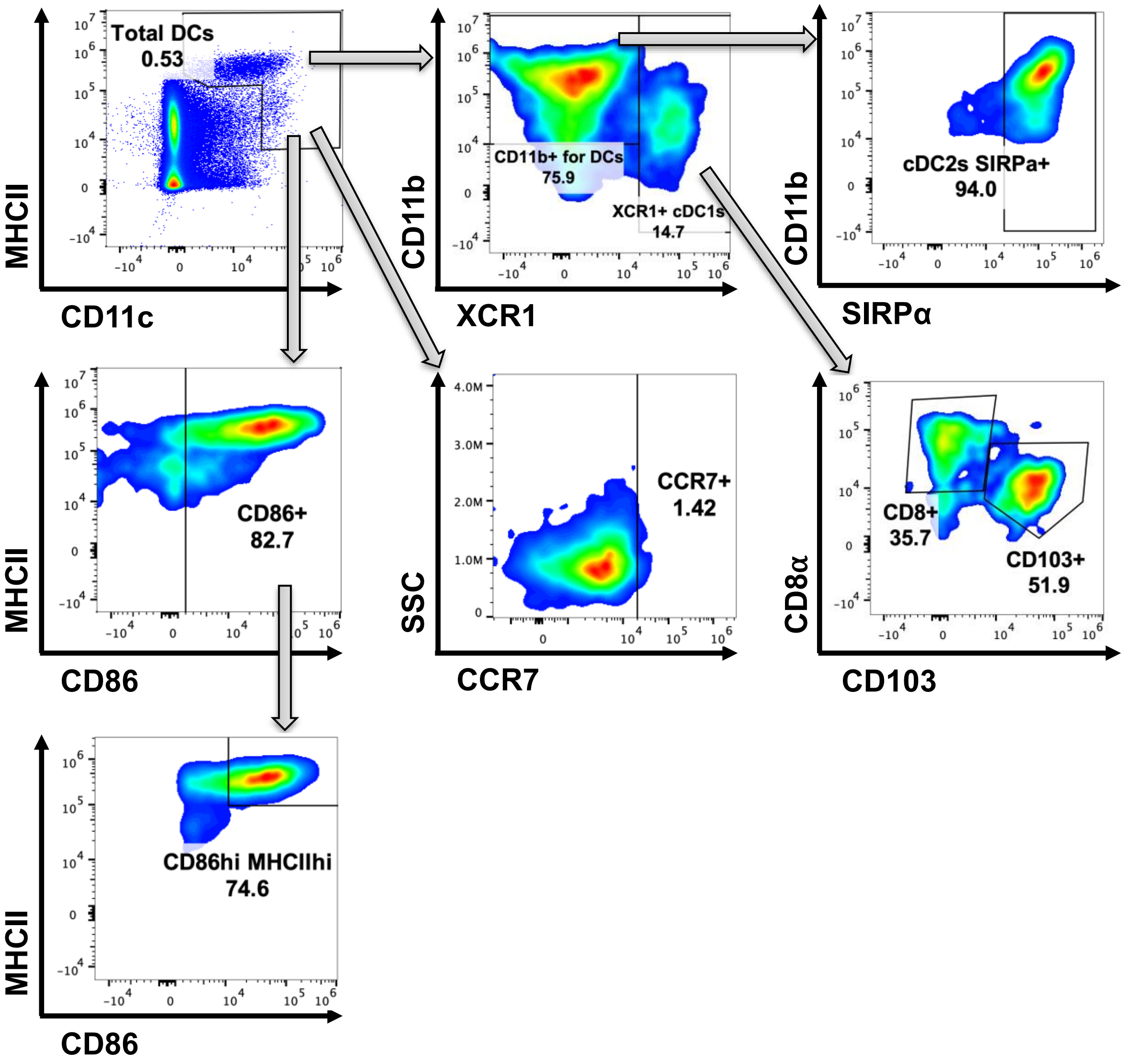


**Figure S7. Gating strategy for flow cytometry analysis of cDCs and activated status.** Gating strategy for the following DC (LiveCD45$^+$CD11c$^+$MHCII$^+$) subsets: cDC1s (XCR1$^+$), migratory cDC1s (XCR1$^+$CD103$^+$), tissue-resident cDC1s (XCR1$^+$CD8α$^+$CD103$^-$) and cDC2s (XCR1$^-$CD11b$^+$SIPRα$^+$). Activation of DC subsets was defined as a subset of CD86$^+$MHCII$^+$: CD86$^{hi}$MHCII$^{hi}$. CD86$^+$ and CCR7$^+$ gating strategies are shown.

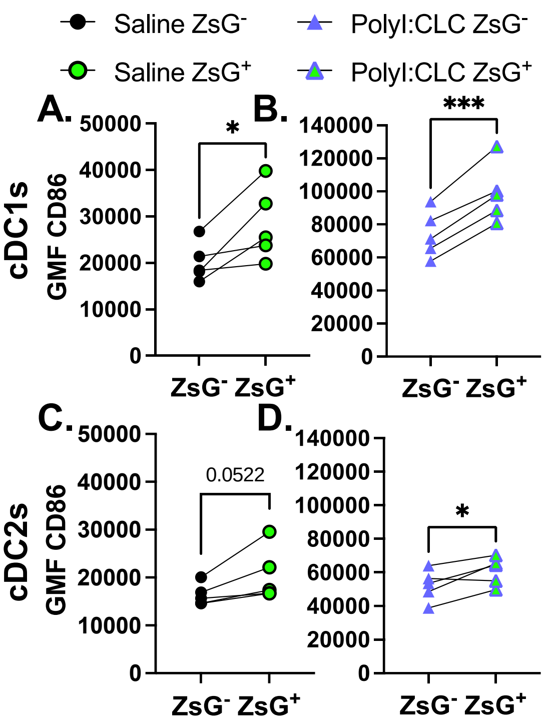


**Figure S8. CD86 expression on cDCs is augmented by toll-like receptor 3 agonist polyI:CLC and ZsG.** 400k B16F10-ZsG cells were inoculated in the right flanks of C57/Bl6 mice and tumors were treated with polyI:CLC (75 mg/0.1 mL saline I.P.) or saline (0.1 mL I.P.) 13 days post inoculation. Axial and brachial lymph nodes were harvested and pooled 24 h after saline or polyI:CLC injection. **A, B)** Geometric mean fluorescent (GMF) intensity of CD86 on cDC1s as a function of ZsG positivity. **C, D)** GMF intensity of CD86 on cDC2s as a function of ZsG. n = 10. Paired t-test: $* p < 0.05$, $** p < 0.01$ , $*** p < 0.001$. Means ± SEM.

**Table S1. Logistics model parameters.**

| Tumor Side | $R^2$ | | $r$ | | $a$ | |
|---|---|---|---|---|---|---|
| | Sham (n = 11) | BH (n = 8) | Sham (n = 11) | BH (n = 8) | Sham (n = 11) | BH (n = 8) |
| **Ipsilateral** | 0.9678 ± 0.0077 (0.9086 to 0.9933) | 0.7993 ± 0.1038 (0.0902 to 0.9956) | $(4.66 \pm 2.58)x10^4$ ($897$ to $2.93x10^5$) | $(3.34 \pm 3.01)x10^6$ ($0.0351$ to $2.44x10^6$) | 0.4626 ± 0.0263 (0.2888 to 0.5908) | 0.4013 ± 0.0431 (0. 2465 to 0.5792) |
| **Contralateral** | 0.9346 ± 0.0429 (0.5127 to 0.9975) | 0.9393 ± 0.0298 (0.7489 to 0.9954) | $(1.43 \pm 0.51)x10^4$ ($0.0094$ to $4.77x10^4$) | $(3.31 \pm 1.25)x10^4$ ($53.55$ to $9.67\ x10^4$) | 0.3625 ± 0.0370 (0.0907 to 0.5028) | 0.3018 ± 0.0497 (0.0250 to 0.4304) |

**Values are Means ± SEM. Parentheses show range, from minimum to maximum. Sample sizes indicated below group labels.2**

**Table S2. Key Resources Table.**

| REAGENTS | SOURCE | IDENTIFIER |
|---|---|---|
| Anti-mouse CD45 clone 30-F11 BUV395 | BD | Cat# 564279; RRID: AB_2651134 |
| Anti-mouse CD3 clone 145-2C11 BUV563 | BD | Cat# 749277; RRID: AB_287352 |
| Anti-mouse CD19 clone 1D3 BUV737 | BD | Cat# 612781; RRID: AB_2870111 |
| Anti-mouse F4/80 clone BM8 BV421 | BioLegend | Cat# 123137; RRID: AB_2563102 |
| Anti-mouse CD11c clone N418 eF450 | Invitrogen | Cat# 48-0114-82; RRID: AB_1548654 |
| Anti-mouse Ly6G clone 1A8 BV605 | BioLegend | Cat# 127639; RRID: AB_2565880 |
| Anti-mouse/rat XCR1 clone ZET BV650 | BioLegend | Cat# 148220; RRID: AB_2566410 |
| Anti-mouse CD103 clone 2E7 BV785 | BioLegend | Cat# 121439; RRID: AB_2800588 |
| Anti-mouse Ly-6C clone HK1.4 PE/Dazzle 594 | BioLegend | Cat# 128044; RRID: AB_2566577 |
| Anti-mouse CD172a (SIRP$\alpha^+$) clone P84 PE | BioLegend | Cat# 144012; RRID: AB_2563550 |
| Anti-mouse/human CD11b clone M1/70 PE/Cy5 | BioLegend | Cat# 101209; RRID: AB_312792 |
| Anti-mouse CD197 (CCR7) clone 4B12 PE/Cy7 | Invitrogen | Cat# 25-1971-82; RRID: AB_469652 |
| Anti-mouse CD86 (B7-2) clone GL1 APC | Invitrogen | Cat# 17-0862-82; RRID: AB_469419 |
| Anti-mouse MHCII clone M5/114.15.2 AF700 | Invitrogen | Cat# 56-5321-80; RRID: AB_494010 |
| Anti-mouse CD8$\alpha$ clone 53-6.7 APC/Fire 750 | BioLegend | Cat# 100766; RRID: AB_2572113 |
| Fixable Live/Dead Blue | Invitrogen | Cat# L23105; UV Excitation |
| DAPI | Sigma-Aldrich | Cat# D9542 |
| Fc Block | Invitrogen | Cat# 14-0161-86; RRID: AB_467135 |
| Brilliant Stain Buffer | BD | Cat# 563794 |
| FACS Lysis | BD | Cat# 349202 |
